# Recent loss of the Dim2 DNA methyltransferase decreases mutation rate in repeats and changes evolutionary trajectory in a fungal pathogen

**DOI:** 10.1101/2020.03.27.012203

**Authors:** Mareike Möller, Michael Habig, Cécile Lorrain, Alice Feurtey, Janine Haueisen, Wagner C. Fagundes, Alireza Alizadeh, Michael Freitag, Eva H. Stukenbrock

**Affiliations:** Environmental Genomics, Christian-Albrechts University, Am Botanischen Garten 1-9, D-24118 Kiel, Germany; Max Planck Institute for Evolutionary Biology, August-Thienemann-Str. 2, D-24306 Plön, Germany; Department of Plant Protection, Faculty of Agriculture, Azarbaijan Shahid Madani University, Tabriz, Iran; Department of Biochemistry and Biophysics, Oregon State University, Corvallis, OR, United States of America

**Keywords:** DNA methylation, genome evolution, transposable elements, experimental evolution, fungal pathogen

## Abstract

DNA methylation is found throughout all domains of life, yet the extent and function of DNA methylation differ between eukaryotes. Strains of the plant pathogenic fungus *Zymoseptoria tritici* appeared to lack cytosine DNA methylation (5mC) because gene amplification followed by Repeat-Induced Point mutation (RIP) resulted in the inactivation of the *dim2* DNA methyltransferase gene. 5mC is, however, present in closely related sister species. We demonstrate that inactivation of *dim2* occurred recently as some *Z. tritici* isolates carry a functional *dim2* gene. Moreover, we show that *dim2* inactivation occurred by a different path than previously hypothesized. We mapped the genome-wide distribution of 5mC in strains with and without functional *dim2*. Presence of functional *dim2* correlates with high levels of 5mC in transposable elements (TEs), suggesting a role in genome defense. We identified low levels of 5mC in strains carrying inactive *dim2* alleles, suggesting that 5mC is maintained over time, presumably by an active Dnmt5 DNA methyltransferase. Integration of a functional *dim2* allele in strains with mutated *dim2* restored normal 5mC levels, demonstrating *de novo* cytosine methylation activity of *dim2*. To assess the importance of 5mC for genome evolution, we performed an evolution experiment, comparing genomes of strains with high levels of 5mC to genomes of strains lacking *dim2*. We found that the presence of *dim2* alters nucleotide composition by promoting C to T transitions (C→T) specifically at CpA (CA) sites during mitosis, likely contributing to TE inactivation. Our results show that 5mC density at TEs is a polymorphic trait in *Z. tritici* populations that can impact genome evolution.

**Author Summary:** Cytosine DNA methylation (5mC) is known to silence transposable elements in fungi and thereby appears to contribute to genome stability. The genomes of plant pathogenic fungi are highly diverse, differing substantially in transposon content and distribution. Here, we show extensive differences of 5mC levels within a single species of an important wheat pathogen. These differences were caused by inactivation of the DNA methyltransferase Dim2 in the majority of studied isolates. Presence of widespread 5mC increased point mutation rates in regions with active or mutated transposable elements during mitosis. The mutation pattern is dependent on the presence of Dim2 and resembles a mitotic version of Repeat-Induced Point mutation (RIP). Thus, loss of 5mC may represent an evolutionary trade-off offering adaptive potential at the cost of transposon control.

## Introduction

DNA methylation is an important process for epigenetic regulation, and functions range from dynamic control of gene expression to transposon silencing and the maintenance of genome integrity [1,2]. Although DNA methylation has been detected on both cytosines and adenines in eukaryotes, cytosine DNA methylation (5mC) has been the focus of most studies so far. In mammals, cytosine methylation is mainly found in “CpGs” (cytosine followed by guanine; here abbreviated CG), while non-CpG methylation (CHG or CHH, where H is any nucleotide other than G) is commonly detected in plants and fungi [3,4]. Various DNA methyltransferases (DNMTs) are involved in the establishment and maintenance of DNA methylation but the distribution and number of enzymes involved is highly variable in different kingdoms [5]. In mammals and plants, enzymes of the DNMT1/MET1 class are maintenance methyltransferases that detect hemi-methylated DNA sequences, for example after replication [6–8]. *De novo* methyltransferases, like DNMT3a and DNMT3b in mammals [9] and DRM in plants [10] act on sequences that are free of methylation, presumably by recognition of specific motifs or patterns [9,11]. Although DNMTs are often classified as maintenance or *de novo* enzymes their function is not necessarily limited to one or the other [12,13].

In fungi, four classes of DNMTs have been identified. Basidiomycetes have DNMT1 homologs that appear to be classical maintenance DNMTs [4,14]. In the ascomycete *Neurospora crassa*, DNA methylation is mediated by a single enzyme, DIM-2 [15]. While the conserved DIM2 class of enzymes shows limited sequence similarity to DNMT1/MET1 maintenance DNMTs, it is a class specific to fungi [16]. Other fungal proteins resembling DNMT1 include *Ascobolus immersus* Masc1 and *N. crassa* RID [17,18], involved in “Methylation Induced Premeiotically” (MIP) or “Repeat-Induced Point mutation” (RIP), respectively [17,19]. DNMT5 enzymes constitute a more recently discovered class of maintenance DNMTs in fungi [20,21]. So far, the presence of 5mC in fungi has been mainly associated with repetitive DNA suggesting a role in genome defense by silencing of transposable elements (TEs). There is little or no evidence that 5mC is also found in coding sequences to influence gene expression [22], although there are examples of promoter methylation [23,24].

A previous study demonstrated amplification of *dim2*, a gene encoding the homolog of *N. crassa* DIM-2, and inactivation by RIP in the genome of the plant pathogenic fungus *Zymoseptoria tritici* [25,26]. For example, the genome of the reference isolate IPO323 carries 23 complete or partial, nonfunctional copies of *dim2*; all alleles show signatures of RIP, namely numerous C:G to T:A transition mutations. Consequently, 5mC was not detected by mass spectrometry in IPO323 [25,26]. However, the amplification of *dim2* must have occurred recently as two closely related sister species of *Z. tritici, Zymoseptoria ardabiliae* and *Zymoseptoria pseudotritici* were shown to carry a single intact *dim2* gene and have 5mC [25].

Population genomic analyses have revealed high levels of genetic variation within and between populations of *Z. tritici* [27–29]. This variation is generated by high mutation and recombination rates and by extensive gene flow. Notably, a dynamic landscape of transposable elements in this fungus has been associated with rapid evolution, not only in *Z. tritici*, but also in sister species [30,31]. Moreover, a recent study demonstrated a pervasive effect of recurrent introgression between closely related species of *Zymoseptoria* [32]. These interspecific hybridization events are evident from the presence of highly diverged alleles that are maintained over evolutionary time within species.

By genome analyses of multiple *Z. tritici* isolates from the center of origin of the pathogen, the Middle East, we discovered several *Z. tritici* isolates with an intact *dim2* gene. Our findings suggest that the loss of *de novo* 5mC not only occurred very recently but is an ongoing process and polymorphic trait in *Z. tritici*. Here, we address the evolution and function of *dim2* in the *Zymoseptoria* species complex. We show that the presence of an intact and functional *dim2* in some *Z. tritici* isolates corresponds to widespread 5mC in TEs. Integration of a wild-type, intact *dim2* allele into a *dim2*-deficient background restores methylation of previously non-methylated regions. Using comparative genomics and experimental evolution we show that loss of *dim2* and thus 5mC affects nucleotide composition of TEs by influencing mutation rates at CA sites.

## Results

### The *dim2* DNA methyltransferase gene is functional in several *Z. tritici* isolates

Based on our identification of a non-truncated *dim2* gene in an Iranian *Z. tritici* isolate, Zt10, we set out to investigate its recent evolution in the *Zymoseptoria* species complex in a much larger collection of genome sequences than previously [25] available. We used BLAST to search for the sequence of *dim2* in 22 high-quality assemblies obtained by SMRT sequencing of *Z. tritici* (Fig 1) [33–36], as well as 17 genomes of *Z. ardabiliae*, and nine genomes of *Z. brevis* (S1 Table), both closely related sister species of *Z. tritici* and considered to be endemic to Iran. We detected a single *dim2* homolog in each of the *Z. ardabiliae* and *Z. brevis* genomes but found multiple mutated and non-functional copies of *dim2* in 17 out of 22 *Z. tritici* genomes (Fig 1). Five *Z. tritici* isolates contained an intact, non-mutated copy of *dim2* in addition to multiple mutated, non-functional copies, except for isolate Zt469, which contained one functional *dim2* gene and no additional copies. We considered *dim2* genes intact and functional when they did not contain pre-mature stop codons or frameshift mutations. Non-functional copies contained numerous mutations including pre-mature stop codons.

**Fig 1.**
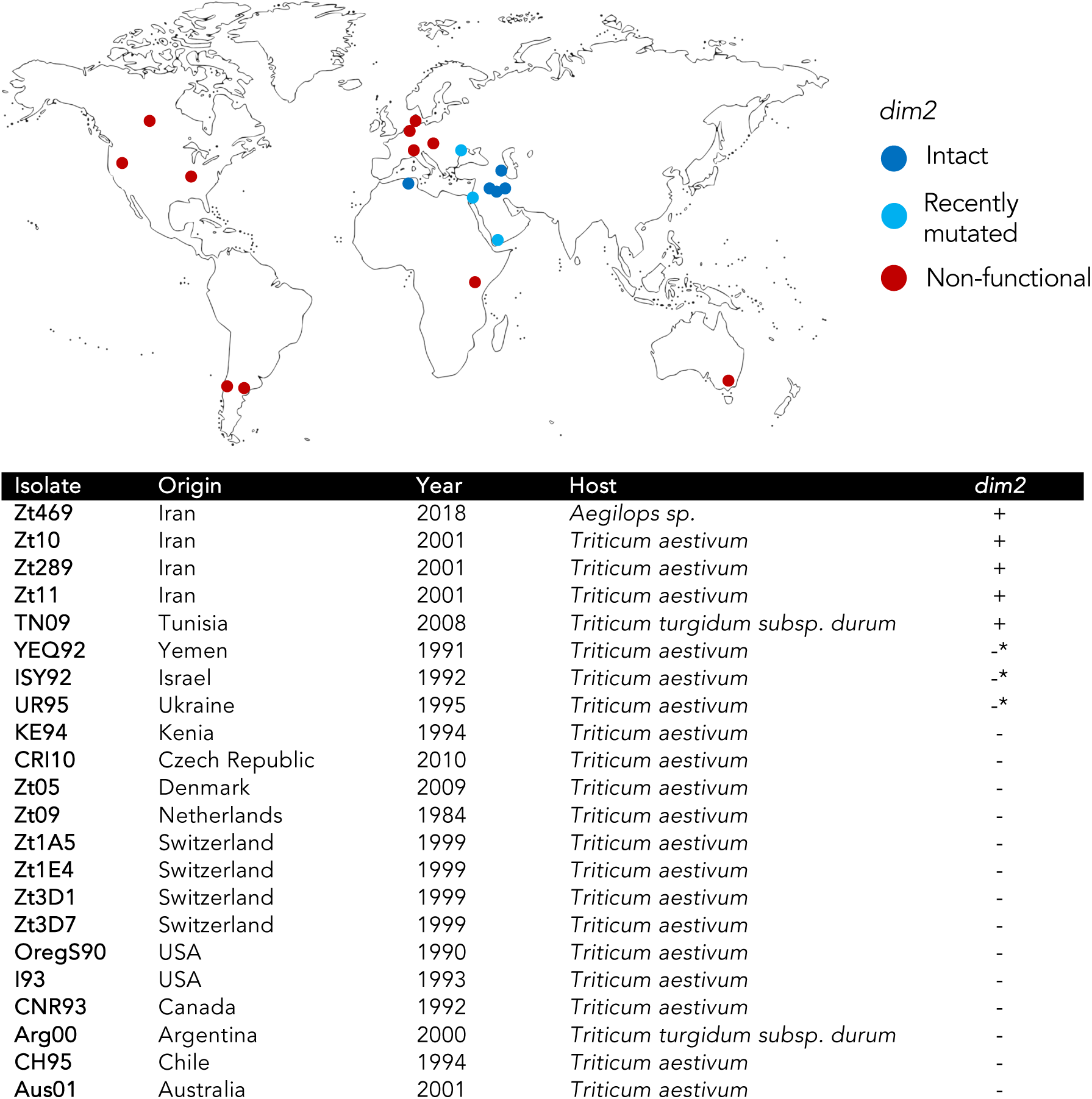
Overview of sampling location, year, host plant, and presence of a functional *dim2* DNA methyltransferase gene of *Z. tritici* isolates analyzed in this study. *Aegilops sp*., wild grass in Triticeae; *Triticum aestivum*, bread wheat; *T. turgidum durum*, Durum wheat. *non-functional, recently mutated native *dim2* gene

Zt469 was isolated from *Aegilops sp*., all other isolates were collected from wheat. Four of those five isolates with intact *dim2* are from Iran (Zt10, Zt11, Zt289, Zt469) [33,37], and one is from Tunisia (TN09, collected from durum wheat) [33]. We further tested whether presence of a non-mutated *dim2* allele is commonly found among Iranian isolates. Twelve additional isolates from Iran (S1 Table) were assayed by PCR and Sanger sequencing and in all cases, we confirmed presence of an intact *dim2* allele. A region of chromosome 6 of IPO323 had been previously identified as the native *dim2* locus; all additional, non-functional copies are located in subtelomeric regions embedded in repetitive DNA [25]. Consistent with this idea, we found that all non-mutated *dim2* genes were located at the predicted native locus on chromosome 6 (Fig 2A, S2 Table).

**Fig 2.**
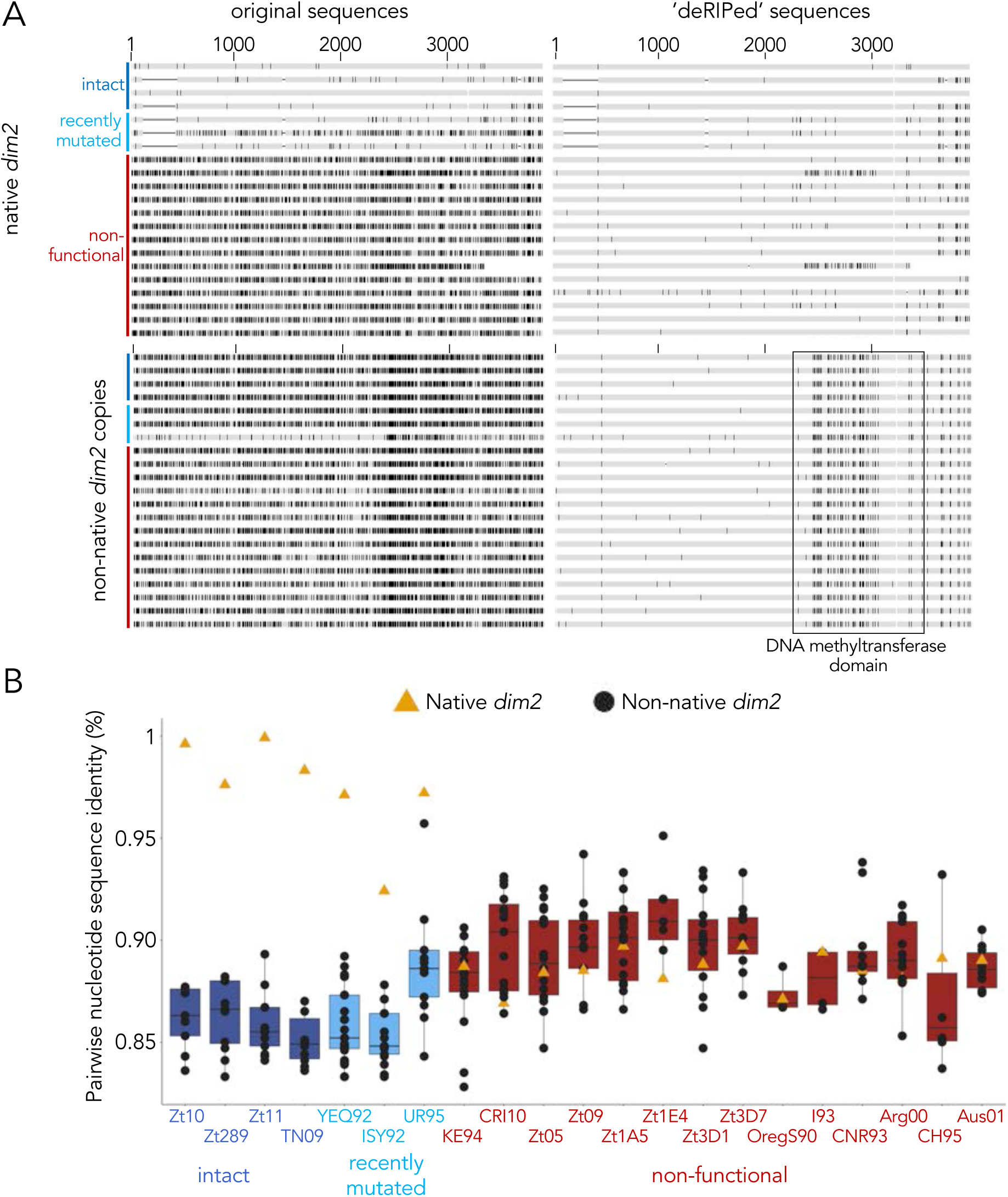
Sequence comparisons of *dim2* alleles in *Z. tritici* isolates. **A)** Mutations of *dim2* in original and ‘deRIPed’ alleles identified by alignment to the functional allele of Zt469. Shown are all native alleles (located on chromosome 6) and one representative non-native, additional copy per isolate. Native alleles (functional and non-functional) lack mutations in the DNA methyltransferase domain that are present in all non-native alleles (black box), except for isolates OregS90 and CRI10. **B)** Pairwise sequence comparison of the native, intact *dim2* allele from Zt469 with all full-length native and additional *dim2* sequences (see S2 Table). No additional *dim2* copies were detected in the genome of Zt469 (from *Aegilops sp*.) whereas all other isolates (from *Triticum* spp.) have additional copies of *dim2*. Non-native, additional copies (black dots) in the isolates with functional or recently mutated *dim2* (except for UR95) showed higher sequence diversity compared to those with inactivated *dim2*, where alleles at the native locus (yellow triangle) are amongst the more diverged copies. This supports the idea that this is the ancestral, now heavily mutated allele of *dim2*. Alleles or isolates are arranged in the same order in panels A (top to bottom) and B (left to right).

### Analyses of *Z. tritici dim2* alleles reveals multiple cycles of recombination and inactivation

Previously, gene amplification of the native *dim2* allele, followed by RIP had been suggested as the most likely cause for the apparent lack of 5mC in the *Z. tritici* IPO323 genome [25,26]. We here conducted a detailed analysis of mutations within native and non-native *dim2* genes in the *Z. tritici* genomes and found some unexpected patterns.

Indeed, as previously reported, non-functional copies of *dim2* contain numerous transition mutations as a consequence of RIP, thus to compare sequences of native and additional (non-native) copies we “deRIPed” mutant copies by reverting C→T and G→A transitions using the functional *dim2* from Zt469 as reference, because it likely represents one ancestral allele. We found that all non-native alleles share a specific pattern of transition and, surprisingly, transversion mutations in the DNA methyltransferase domain that are not found in the vast majority of native *dim2* genes (Figs 2A, S1A and S2). Transversions are not a consequence of RIP [38,39], thus a mixture of transitions, transversions, and deletions suggests that *dim2* was mutated not simply as a result of amplification of the native gene followed by widespread RIP, as originally proposed [25]. Rather, integration of a different *dim2* allele (hereafter called transversion allele) of unknown origin into subtelomeric regions is likely to have resulted in subsequent gene amplification, RIP and presence of multiple non-functional, non-native *dim2* copies. In support of this hypothesis, we found that the flanking regions of the additional, but not the native, copies share homology and are annotated as retrotransposons, suggesting that integration and amplification of non-native copies may have been TE-mediated.

While this idea explains events at subtelomeric integrations sites, we also identified two isolates in which the native *dim2* gene on chromosome 6 shares the specific transversion mutations in the DNA methyltransferase domain, OregS90 and CRI10 (from the USA and Czech Republic, respectively) (Fig 2A). The native OregS90 *dim2* is truncated and surrounded by TEs and the native *dim2* of CRI10 lacks mutations in the 3’ region that are present in all additional, but not native, copies.

Five native *dim2* genes have a shared deletion in the 5’ region that is not found in any of the additional copies. Given the previously identified signatures of introgression in *Z. tritici* genomes we consider that these different *dim2* alleles at the native locus may reflect separate integration events that have been directed by sequence identity. In summary, our analyses revealed that gene amplification followed by RIP alone is insufficient to explain the distribution of variant *dim2* alleles in *Z. tritici* isolates.

To further investigate the evolutionary history of *dim2* in *Z. tritici* isolates, we compared the sequence identity of the non-mutated *dim2* gene from Zt469 to all other, native and additional, copies of *dim2* in the 22 *Z. tritici* genomes (Fig 2B). Gene amplification followed by RIP predicts that by accumulating transition mutations all copies should become quite similar to each other; RIP becomes less efficient or stops when sequence identity between alleles drops to ∼80% [38]. Instead, our analyses revealed that 18/21 isolates showed two distinct patterns, the first matching the prediction because isolates lacking functional *dim2* showed higher levels of DNA sequence identities (85-95%) between native and additional alleles (Fig 2B, S2 Table; red, “non-functional”, pattern 1). Here, all copies have suffered repeated RIP cycles, and all copies are non-functional. When, however, an intact, native *dim2* allele is present (pattern 2), the additional copies show lower DNA sequence identities (84-88%) while the functional alleles are almost identical (>96%) to the Zt469 allele (Fig 2B, S2 Table; dark blue, “intact”; Zt10, Zt289, Zt11, TN09). This second pattern was unexpected; the most parsimonious explanation is replacement of non-functional alleles at the native locus with novel *dim2* alleles, perhaps by recombination with wild grass infecting isolates, such as Zt469.

We discerned a third group of three isolates, which suggests that repeated cycles of transposon- or recombination-mediated amplification of *dim2* alleles are still ongoing (Fig 2B, S2 Table; light blue, “recently inactivated”; YEQ92, ISY92, UR95). YEQ92 and ISY92 contain non-functional *dim2* alleles at the native locus that are still similar (96 and 92%, respectively) to the functional *dim2*, suggesting a more recent inactivation of *dim2* in these isolates. All additional, subtelomeric copies have low sequence identities as in the second pattern described above. UR95 has two recently mutated *dim2* alleles, one at the native locus (Zt469 allele) and one in the subtelomeric region on chromosome 8 (transversion allele); both are non-functional alleles but contain relatively few mutations and are still similar to the functional *dim2* allele (>90%) suggesting recent integration and inactivation. In summary, our findings suggest that there are multiple functional alleles of *dim2* at the native locus present in the *Z. tritici* population. One group of alleles is derived from the Zt469 allele, a second group contains the “fingerprint” of the transversion allele when compared to the Zt469 allele.

### Integration of an intact *dim2* allele at the native locus restores 5mC in Zt09

To assess the role of *dim2* in DNA methylation we generated mutants in which we either restored or deleted *dim2*. We integrated the functional *dim2* gene of the Iranian Zt10 isolate into the native *dim2* locus of Zt09 (derived from reference isolate IPO323) by selecting for hygromycin resistance, conferred by the *hph* gene, generating the strain Zt09::*dim2*. In Zt10, we replaced the functional *dim2* gene with *hph*, generating the strain Zt10Δ*dim2*. Integration or deletion of the *dim2* alleles were verified by Southern analyses (S3 Fig). To test whether *dim2* affects growth or infection, we compared the mutant and respective wild-type strains under different *in vitro* conditions as well as during host infection. We did not detect any differences between wild-type and mutant strains *in vitro* (S4 Fig) but integration of the functional *dim2* in Zt09 significantly reduced necrosis and asexual fruiting body formation (measured as number of asexual fruiting bodies, pycnidia, per leaf area) *in planta* (S5 Fig). We therefore conclude that presence of *dim2* is dispensable under the tested *in vitro* conditions but that re-integration of *dim2* into the Zt09 background decreases virulence *in planta*.

To validate the presence of 5mC and to compare patterns between isolates and mutants, we performed whole-genome bisulfite sequencing (WGBS). We sequenced three replicates of each wild-type isolate (Zt09 and Zt10) and mutant strain (Zt09::*dim2* and Zt10Δ*dim2*). As a control for the bisulfite conversion rate, we added Lambda DNA to each sample; based on this, the bisulfite conversion rate was >99.5%. To confirm the WGBS sequencing results, we performed an additional independent bisulfite conversion, amplified target regions by PCR, cloned and sequenced the fragments, and we performed Southern blot analyses using 5mC-sensitive and -insensitive restriction enzymes (S6 Fig).

We detected high numbers of methylated cytosines in isolates with functional *dim2* alleles, i.e., the wild-type Iranian isolate Zt10 and Zt09::*dim2*. Surprisingly, we also detected low levels of 5mC in the absence of *dim2*, i.e., in Zt10Δ*dim2* and Zt09, which had been missed by the mass spectrometry in earlier studies [25]. While on average ∼300,000 sites in Zt10 and ∼400,000 sites in Zt09::*dim2* showed 5mC in the presence of *dim2*, only ∼12,000 (Zt09) and ∼40,000 (Zt10Δ*dim2*) 5mC sites were found when *dim2* was absent (Fig 3A). In all strains, 5mC was restricted to previously annotated TEs and adjacent non-coding regions (Fig 3A). We did not find evidence for gene body or promoter methylation, except for few TE-derived genes. By correlating the 5mC data with previously published histone methylation maps [40] we found that 5mC co-localizes with H3K9me3 marks (Fig 3B). This finding is consistent with observations in *N. crassa, C. neoformans*, and *A. thaliana* [21,41–44].

**Fig 3.**
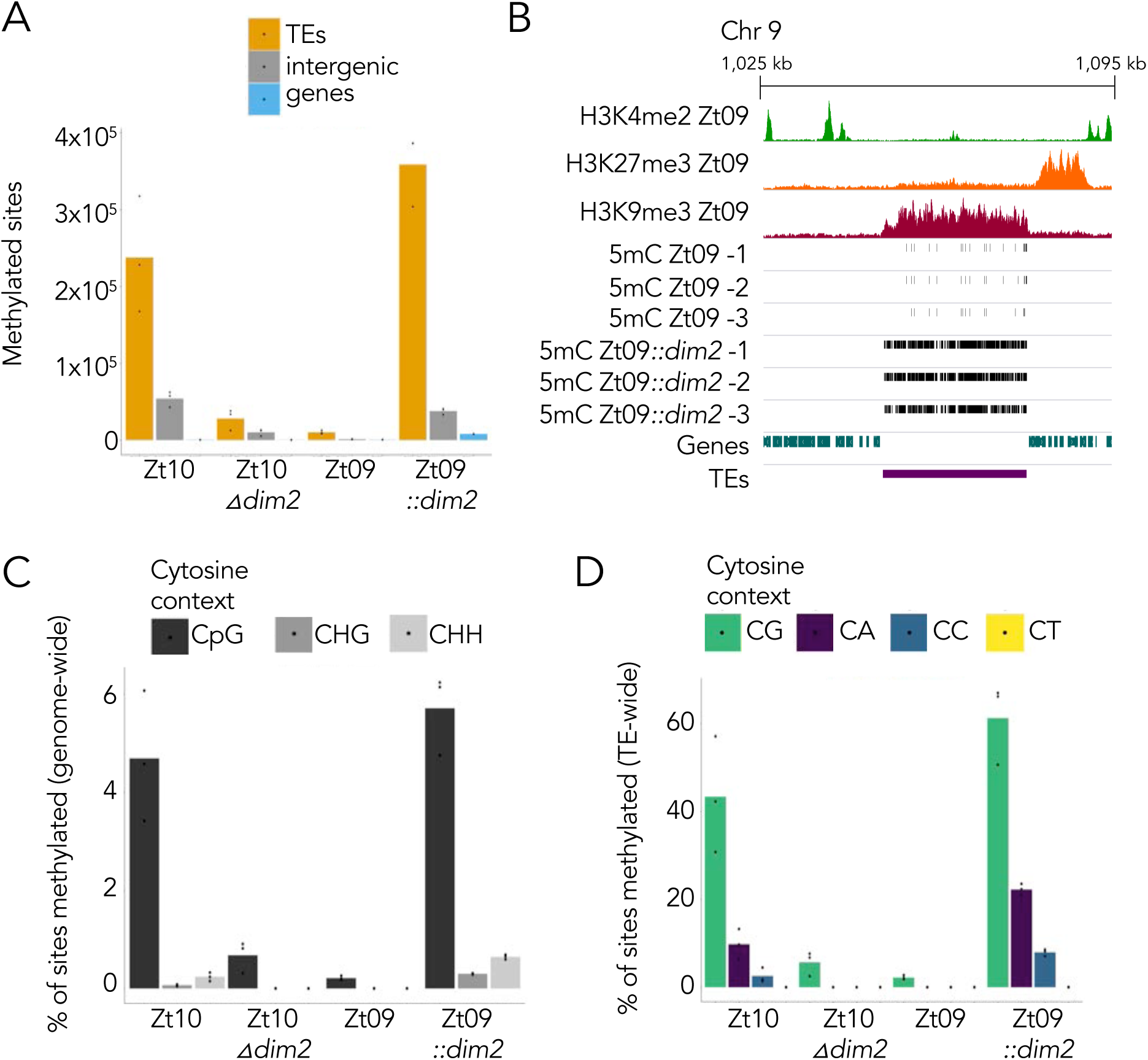
Localization and site preferences of cytosine methylation in *Z. tritici* genomes in presence and absence of *dim2*. **A)** Number of 5mC methylated sites detected by whole genome bisulfite sequencing (WGBS) in TEs, intergenic regions, or genes. The vast majority of methylated sites is localized in TEs followed by intergenic regions and TE-derived genes. **B)** 5mC co-localizes with H3K9me3 and TEs in Zt09. Shown are ChIP-seq tracks for H3K4me2, H3K27me3 and H3K9me3 [40], and 5mC sites detected in a representative region on chromosome 9 in the genomes of Zt09 and Zt09::*dim2* (three replicates). **C)** Genome-wide 5mC levels in the different isolates and respective mutants. 5mC levels are higher in the presence of *dim2*, and *dim2* is required for all non-CG methylation. **D)** 5mC levels and site preferences in TEs. CG sites are preferred, followed by CA, and CC. 5mC is almost completely absent from CTs.

We next analyzed the specific context of DNA methylation. In strains without functional *dim2*, we found 5mC exclusively in CG contexts (genome wide ∼0.2% in Zt09 and ∼1% in Zt10Δ*dim2*; percentage based on all Cs in the genome). In the presence of *dim2*, cytosines in CG as well as CHG and CHH contexts were methylated. Genome-wide, ∼5 or 6% of CGs and <1% of CHGs or CHHs were methylated in Zt10 or Zt09::*dim2*, respectively (Fig 3C). In sequences annotated as TEs, however, up to 60% of all CGs were methylated (Fig 3D). Methylation of CGs occurred most frequently (∼43%, ∼61%), but we also detected methylation at CAs (∼10%, ∼22%), and to lesser extent at CCs (∼3%, ∼8%). At CTs, there was hardly any detectable methylation (<0.01%; Figs 3D, S7) in Zt10 or Zt09::*dim2*, respectively. This was surprising, as CTs are the most abundant CH dinucleotides in TEs while CAs are the least abundant sites in genomes of both Zt09 and Zt10 (S3 Table). This suggests a strong site-specific preference of Dim2 for CAs but not CTs in *Z. tritici*.

Upon integrating of a functional *dim2* allele in Zt09 we found that overall 5mC levels are higher when compared to Zt10 (compare Fig 3C, left- and right-most panels), and CG methylation levels in Zt10Δ*dim2* were higher compared to Zt09 (compare Fig 3C, two middle panels). These findings suggest that 5mC levels can decrease significantly over relatively short evolutionary time spans even if a *de novo* DNA methyltransferase is present.

### DNA methylation can be maintained in absence of *dim2*

We next asked how 5mC can persist in the genome of *Z. tritici* in the absence of a functional *dim2* gene. The presence of CG methylation in isolates without *dim2* suggested the presence of an additional DNMT. Thus, we searched for putative DNMT coding regions in the genomes of *Z. tritici, Z. ardabiliae*, and *Z. brevis* with the conserved DNMT domain of Dim2 as query in a BLASTp search. In all genomes, we found two additional predicted DNMTs (S8 Fig). One is similar to Dnmt5 of *Cryptococcus neoformans* [20] (*dnmt5*, Zt_chr6_00685), the other is a homolog of *N. crassa* RID [17] (*rid*, Zt_chr5_00047; manually corrected gene annotation) [45]. Genes for Dnmt5 and Rid are present in all *Zymoseptoria* spp. genomes we analyzed. The *dnmt5* alleles are highly conserved and show very little inter- or intraspecies diversity (S1B Fig). In contrast, *rid* shows an exceptionally high inter- and intraspecies diversity with three highly distinct alleles present in *Zymoseptoria* spp. (S1C Fig). This diversity may stem from introgression events between *Zymoseptoria* species as recently described for highly variable regions in the genome of *Z. tritici* [32]. However, presence of the different *rid* alleles does not correlate with presence of a functional *dim2* gene in *Z. tritici*.

Masc1 and RID are involved in genome defense during pre-meiosis [17,18]. Although Masc1 is presumed to be a *de novo* DNMT acting during the sexual cycle of *Ascobolus immersus*, there are still no enzyme activity data available for either Masc1 or RID. In contrast, Dnmt5 has been characterized as a 5mCG maintenance DNMT in *C. neoformans* [20,21] and is thus the most likely candidate for maintenance of 5mC at CG sites that we observed here. Further studies will address the role of *dnmt5* in maintenance and impact on 5mC in *Z. tritici*.

### Presence of *dim2* impacts nucleotide composition and activity of TEs

DNA cytosine methylation is correlated with increased C→T mutation rates [46,47]. In *Z. tritici*, and most other fungi [22], 5mC is enriched in TE sequences. To evaluate whether the presence of *dim2* impacts sequence composition of TEs, we analyzed TE sequences in isolates with and without a functional *dim2* gene. Therefore, we extracted TE sequences, calculated dinucleotide frequencies relative to the TE content for each genome (S3 Table), and visualized the difference in frequencies by correspondence analysis. Isolates lacking functional *dim2* have more CA/TG sites within TEs compared to isolates with functional *dim2* (Fig 4A, B). Conversely, isolates with functional *dim2* contain more TA sites in TEs (Fig 4A, B). Two isolates with recently inactivated *dim2*, ISY92 and YEQ92, show the same pattern as isolates with functional *dim2* but UR95, with two more recently inactivated copies, groups with isolates lacking functional *dim2*. CA/TG sites are the least frequent sites in TEs, while TA sites are the most abundant. We observed the same pattern (i.e., increased number of TA sites), when we included the sister species *Z. ardabiliae* (Za17) and *Z. brevis* (Zb87), and the more distantly related barley pathogen, *Z. passerinii* (Zpa63), all of which contain a single, functional *dim2* (Fig 4B; S3 Table). This pattern is only detectable in TEs, not in the rest of the genome (S3 Table).

**Fig 4.**
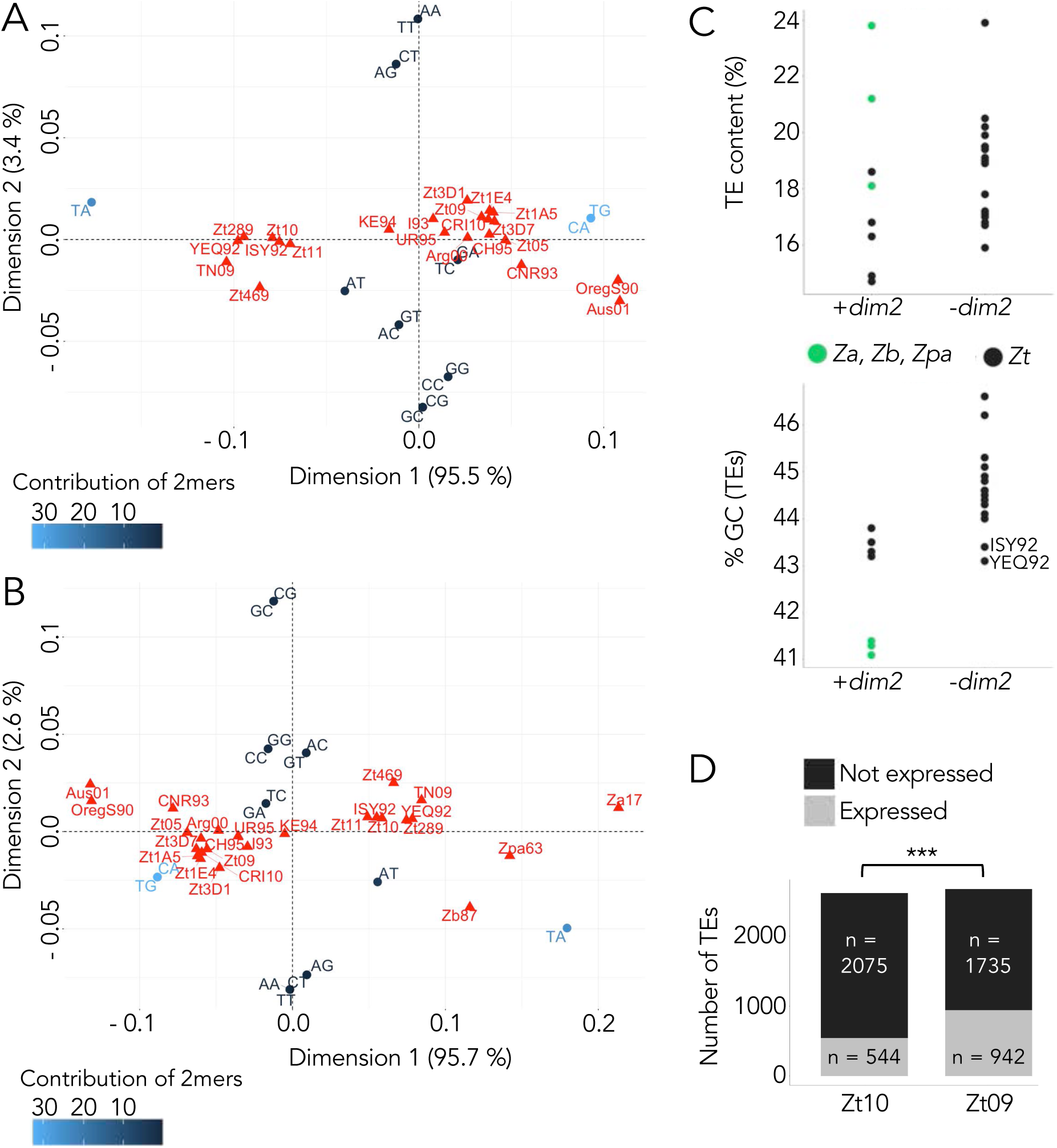
Impact of Dim2 on nucleotide composition and TE activity. **A)** and **B)** Correspondence analyses showing differences in dinucleotide (2mer) frequencies in TEs of different *Z. tritici* isolates **(A)** and other members of the *Zymoseptoria* species complex **(B)**. TA and CA/TG dinucleotide frequencies have the highest impact on the observed differences between isolates and species, indicated by their relative positions on the dimension 1 *versus* dimension 2 plot. Isolates (even from different species) containing an intact *dim2* (Zt10, Zt289, Zt11, Zt469, TN09, Zb87, Za17, Zpa63) and recently mutated *dim2* (ISY92 and YEQ92) have higher TA content and cluster together, while isolates without functional *dim2* have higher CA/TG frequencies in TE sequences. **C)** Comparison of TE and GC content between different *Z. tritici* isolates and *Zymoseptoria* species with and without functional *dim2*. **D)** Number of expressed TEs during the time course of wheat infection is significantly higher in *Z. tritici* isolate Zt09 (non-functional *dim2*) compared to Zt10 (intact *dim2*) (Fisher’s Exact Test for Count Data, *p-*value < 2.2 x 10^−16^).

Accelerated mutation rates in TEs may impact TE activity, so we next determined the TE content of the various *Z. tritici* isolates and compared it to that observed in other *Zymoseptoria* species (Fig 4C). Isolates with functional *dim2* had slightly lower TE content than those lacking *dim2* (∼16.2% *vs*. ∼18.7%; Fig 4C) but TE content in other *Zymoseptoria* species, all with functional *dim2*, was higher than in many *Z. tritici* strains. Thus, there is no simple or clear correlation between TE content and *de novo* 5mC activity. GC content of TEs, however, is considerably lower in isolates with functional *dim2* and two isolates with recently mutated *dim2*, ISY92 and YEQ92, further indicating that Dim2 impacts nucleotide composition (Fig 4C).

The TE content of a genome does not necessarily reflect the activity of TEs, as these sequences may be relics, i.e., mutated and thus inactivated TEs. We therefore assessed whether TEs in Zt09 and Zt10 contained annotated, transposon-related genes as a putative indicator for the presence of active TEs. In Zt09, 56 genes completely overlap (>90% of the sequence) with annotated TEs. Among those, we identified 30 transposon- or virus-related genes encoding for endonucleases, transposases, ribonucleases, and reverse transcriptases, or genes encoding virus-related domains (S4 Table). In contrast, we did not detect any fungal genes overlapping TEs in Zt10. We did find homologs of some of the transposon-associated genes of Zt09 in the genome of Zt10, but they all contained numerous transitions resulting in mis- and nonsense mutations. This indicates that Zt09 contains active TEs whereas homologous elements are inactivated in Zt10.

Lastly, we compared the expression of TEs in Zt09 and Zt10 during the time course of infection by analyzing previously published RNA-seq data [36]. We computed transcripts per million (TPM) originating from TEs in both the biotrophic and necrotrophic phases of infection and considered a TE expressed when TPM was > 0. While ∼21% of TEs produced RNA in Zt10, ∼35% of TEs were transcribed in Zt09 (Fig 4D) indicating that silencing of TE loci is less pronounced in Zt09 where *dim2* is non-functional.

### Dim2 promotes C→T transitions at CAs during mitosis

The difference between CA/TG and TA site abundance in genomes with and without functional *dim2* suggests that the mutation rate, specifically for C→T transitions, is affected by the presence of 5mC. To address whether Dim2 plays a role in promoting these mutations, we conducted a one-year evolution experiment that included the Iranian isolate Zt10 (functional *dim2*) and the reference isolate IPO323 (non-functional *dim2*, isolate Zt09 is a derivate of IPO323). This was a mutation accumulation experiment without competition or selection on cells that were dividing by mitosis only. Each ancestral isolate was propagated as 40 independent replicates. After 52 weeks (corresponding to ∼1,000 mitotic cell divisions) the genomes of the 40 evolved replicate strains per isolate and the progenitor isolates were sequenced to map SNPs. We found that the mutation rate in Zt10 was more than tenfold higher than in IPO323, on average ∼170 *vs*. ∼13 mutations per replicate (Fig 5A, S5 Table). The vast majority of mutations in Zt10 (> 95%) occurred in 5mC regions (S6 Table) and > 95% of all mutations were C→T transitions, while only ∼32% of all mutations in IPO323 were C→T transitions. In Zt10, > 98% of the C→T transitions occurred at CAs (*vs*. ∼21% in IPO323) (Fig 5B). This was a surprise, because the majority of methylated sites are at CGs. Our findings suggest the presence of a *dim2*-dependent mechanism that specifically targets and mutates CA/TG but not CGs in sequence repeats during mitosis. Such a mutator resembles the hitherto undescribed but much looked-for mitotic version of RIP.

**Fig 5.**
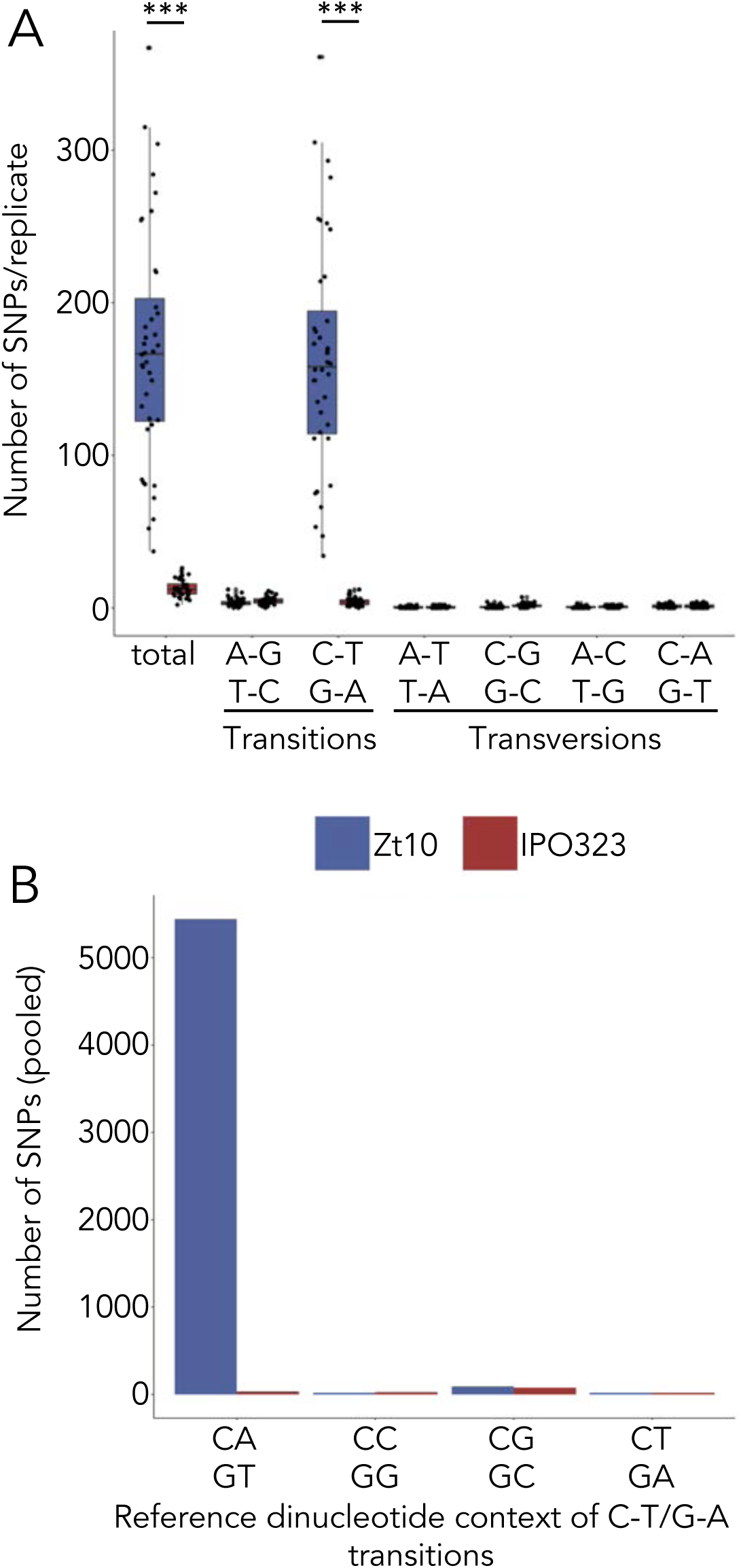
Number and site preferences of SNPs in isolates with functional (Zt10) and non-functional (IPO323*) *dim2* after one year of mitotic growth. **A)** Detected number of SNPs in the 40 replicates of each isolate. The number of SNPs in Zt10 replicates is significantly higher compared to IPO323 (Wilcoxon rank sum test, \*\*\**p-*value < 0.001). The vast majority of mutations are C to T transitions. **B)** Sequence context of the mutations in the two tested isolates. The SNPs of all replicates were pooled for this analysis. C→T transitions are predominantly found in a CA context resulting in CA to TA (or GT to TA) mutations. CG sites, although preferentially methylated, are not mutated at high rates. *Isolate Zt09 is a derivate of IPO323.

## Discussion

Our study of the DNA methyltransferase *dim2* in the fungus *Z. tritici* reveals an exceptional extent of intra species polymorphism that has functional and genetic implications. Although it has been reported that 5mC is absent in *Z. tritici* [25], we found that some isolates contain a functional Dim2 *de novo* 5mC methyltransferase. We show that loss of most 5mC is quite recent; we found that 5mC is not completely absent in *Z. tritici*, even when there is no functional Dim2; and we identified a Dim2-dependent mutator phenomenon. The majority of isolates in which we identified non-mutated *dim2* genes originated in Iran, previously shown to be the center of origin of *Z. tritici* [37]. The closely related sister species *Z. ardabiliae* and *Z. brevis* were collected in Iran as well and are so far considered to be endemic to this region [48,49]. We found only one *Z. tritici* isolate (Zt469) that did not carry any additional *dim2* copies. This isolate was collected from *Aegilops* sp., also in Iran. We propose that the original state of the native *dim2* locus has been maintained in the two sister species and *Z. tritici* isolates collected from wild grasses. Even functional *dim2* alleles show sequence polymorphisms, i.e., transitions and mid-size in-frame deletions, respectively. Thus, all available data suggest that *dim2* gene amplification, recombination events, and RIP have been specific to wheat-infecting *Z. tritici* isolates.

Here, we disproved a previous hypothesis, suggesting that the mutated *dim2* copies in IPO323 and similar strains arose by gene duplications of the native *dim2* followed by transposon-mediated movement to subtelomeric regions and RIP [25]. Instead, DNA sequences of the non-functional subtelomeric *dim2* copies show patterns of increased transversion mutations specifically in the DNMT catalytic domain of the gene. This pattern is not an expected outcome of RIP. Instead we consider interspecific hybridization and introgression between distinct lineages with different *dim2* alleles. Our findings suggest that there are at least two groups of functional *dim2* alleles at the native locus in the extant *Z. tritici* population. One group is derived from or similar to the Zt469 allele, a second group of alleles is enriched with “transversions” when compared to the Zt469 allele. Two strains, OregS90 and CRI10, had non-functional “transversion alleles” at the native locus and we speculate that both may be derived by secondary integration events. Isolate UR95 contains numerous non-functional copies but also two more recently mutated alleles, one Zt469-type and one transversion allele, both non-functional. This indicates that intact and functional copies of both types of alleles are still present in *Zymoseptoria* populations. The current set of genomes did not reveal a functional “transversion allele” at the native locus or subtelomeric region, but we identified a putative functional copy in WAI329, an Australian Illumina-sequenced isolate [50]. A likely scenario for repeated invasion and inactivation cycles predicts that some wild grass infecting isolates carried “transversion alleles” that were not mutated. Gene amplification or TE-mediated recombination coupled to RIP resulted in numerous non-functional alleles. Our data suggest recombination between isolates with *dim2* non-native transversion alleles, which may have induced RIP in the native *dim2* allele because of high sequence similarity outside of the DNMT catalytic domain, thus resulting in non-functional *dim2* alleles only, and therefore absence of *de novo* DNMT activity. This is consistent with the previously reported signatures of re-current introgression in *Z. tritici* [32]. Further analyses of diverse *Z. tritici* isolates from different hosts and geographical locations (*T. aestivum, Triticum durum*, or *Aegilops* sp.) are needed to track the evolutionary history of the *dim2* gene in the *Zymoseptoria* species complex.

The previous study on *dim2* suggested absence of 5mC based on liquid chromatography coupled with electrospray ionization tandem mass spectrometry (ESI–MS/MS) [25]. This finding is consistent with the low levels of methylation we discovered in strains lacking Dim2 activity. Based on recent results obtained in studies with *Cryptococcus* we hypothesize that this low level of 5mC has been maintained by the single homologue of Dnmt5 [20,21] after the loss of Dim2. This DNMT acts as a maintenance DNMT but does not catalyze *de novo* DNA methylation [21]. In the absence of a *de novo* DNMT, DNA methylation is likely to decrease over time as any loss of 5mC cannot be restored. This idea is supported by our finding that 5mC levels are higher upon deletion of *dim2* (Zt10Δ*dim2*) compared to Zt09, where *dim2* was likely inactivated thousands of generations ago.

Sequence repeats are the predominant targets of DNA methylation in fungi [15,22,51,52], and 5mC enrichment is correlated with genome defense mechanisms that act to silence transposons [1]. We found that nucleotide composition in TEs differs between isolates and species that have functional or non-functional *dim2* alleles. CA sites, the preferred target sites of Dim2 in addition to CGs, are reduced in frequency when Dim2 is present. TA sites, however, are more frequent when Dim2 is present. This links increased C→T transition frequency to the presence of functional Dim2. Spontaneous deamination of 5mC also yields C→T transitions [47], giving rise to accelerated mutation rates at 5mC sites, a well-known phenomenon [53]. Correlation of decreased C:G to T:G transitions in absence of the DNA methyltransferase *CMT3* was recently shown in plants [54]. However, the observed differences in dinucleotide frequencies in *Zymoseptoria* and the high rate of C→T transitions in the evolution experiment suggest a role of Dim2 and 5mC in the generation of mutations. The mutations occurring throughout the evolution experiment specifically affected CAs but not CGs. If mutations resulted from spontaneous deamination, we would have expected a greater effect on CGs because they have higher 5mC levels. Lack of Dim2 resulted in complete loss of 5mC at CAs and therefore reduced CA to TA mutation frequencies. We also did not observe a noticeable difference of CG site abundance between isolates with or without functional Dim2, suggesting that non-CG 5mC (mediated by Dim2) is the main driver of the C→T transitions we observed. This suggests that, although repetitive regions are a shared target for DNMTs in different fungal species, distinct target sites of 5mC may correlate with the sequence composition of repetitive regions and the presence of other proteins involved in the 5mC pathway.

What is the underlying mechanism for the reduced CA to TA mutation frequency in absence of functional *dim2*? The best-known process causing C→T transitions in fungi is RIP, which occurs during pre-meiosis after fertilization but before karyogamy [38,39]. While RIP depends largely on the presence of the putative DNMT, RID [17], DIM-2 has also been suggested to be involved in RIP in *N. crassa* [19]. Thus, we propose that the absence of Dim2 may also play a role in the efficiency and rate of RIP in *Z. tritici*. Mechanistic studies are lacking to address this question.

There is little precedence for a mutator process that generates CA to TA mutations and acts during vegetative growth in any eukaryote, the type of mutator uncovered by our evolution experiment. In *N. crassa*, RIP acts preferentially on CAs [19,39,55], and DIM-2 preferentially methylates CTs [22]. Presence of *Z. tritici* Dim2 yields non-CpG 5mC preferentially at CAs and increases C→T mutations. Essentially, the *dim2*-dependent phenomenon described here resembles what is expected for a mitotic version of RIP. We do not know yet whether Dim2 is directly involved in this mechanism and whether Rid may also play a role. However, functional alleles of *rid* are present in all strains we analyzed in the evolution experiment, and in all species examined so far *rid* is expressed solely late during the sexual cycle [17]. Experiments with bacterial DNMTs uncovered inherent mutator activity under appropriate environmental conditions (e.g., low concentrations of the methyl donor, *S*-adenosylmethionine [SAM]) or when specific point mutations (e.g., in the SAM-binding motif) where generated [56–61]. Another possible mechanism may involve differences in DNA repair efficiency where CG sites are more efficiently repaired compared to CA sites. Hemi-methylation of mutated CG sites could hereby aid in strand recognition and mismatch repair as suggested in mammalian cells [62]. Our ongoing studies are directed towards uncovering sequence motifs in Dim2 and examining growth conditions that increase or decrease the novel mutator phenotype.

Why are presence of widespread 5mC and differences in mutation rates polymorphic traits in this fungal pathogen? In several fungal species, 5mC is completely absent or methylation levels are very low [22,63–65] raising the question about the importance of 5mC in terms of genome defense and evolution. While silencing or inactivation of TEs protects genome integrity and stability, it may limit the adaptive potential in response to changing environmental conditions. Transposon-mediated genome diversity, gene or even chromosome copy number variations are frequently found in fungal pathogens, and they form a crucial aspect of rapid evolution [66–69]. Here we uncovered a system of continuing loss and re-acquisition of the ability to silence or inactivate TEs in the genomes of numerous isolates of an important plant pathogen, *Z. tritici*. Based on currently available data, increased 5mC levels negatively affect virulence on wheat, suggesting that loss of *dim2* is adaptive for survival in agricultural environments. Nevertheless, in the center of origin, functional *dim2* copies are re-acquired, suggesting that 5mC is important under certain environmental conditions. The efficiency and abundance of DNA methylation and mutations may therefore represent an evolutionary trade-off between genome integrity and adaptive potential.

## Materials and Methods

### Fungal isolates and growth conditions

All *Zymoseptoria* spp. isolates used in this study (Table S1) were cultivated at 18°C in YMS (4 g yeast extract, 4 g malt, 4 g sucrose per 1 L, 20 g agar per L for plates) medium. Cultures for DNA extraction and plant infection experiments were inoculated directly from the −80°C glycerol stocks and grown in liquid YMS medium at 200 rpm for 5 days (DNA extractions) and in pre-cultures (3 days) and main cultures (2 days) for plant infections.

### SMRT sequencing and assembly of Iranian isolates Zt289 and Zt469

High molecular weight DNA was extracted as previously described [70]. Library preparation and PacBio sequencing was performed at the Max Planck Genome Center in Cologne, Germany (https://mpgc.mpipz.mpg.de/home/) with a Pacific Biosciences Sequel II. One SMRT cell was sequenced per genome. Genome assemblies were performed as described [71]. Genome assembly statistics are summarized in Table S7.

### Identification of TEs and analysis of TE expression

We annotated transposable elements with the REPET pipeline (https://urgi.versailles.inra.fr/Tools/REPET) [72] as described in [71]. Briefly, we identified repetitive element consensus sequences in each genome using TEdenovo following the developer’s recommendations [72]. We used each library of consensus sequences to annotate genomes using TEannot with default parameters. To evaluate TE activity we analyzed expression *in planta* using RNA-seq data of *Z. tritici* isolates Zt10 and Zt09 [36]. First, raw sequencing reads were quality filtered and trimmed using Trimmomatic [73] and mapped to the respective genome using hisat2 [74]. Reads were trimmed using the following parameters: LEADING:30 SLIDINGWINDOW:4:30 AVGQUAL:30 MINLEN:50. After mapping, the resulting aligned read files were used to assess read counts for each stage and replicate (with a total of two replicates per stage). The read count table was generated using TEcount function of the TEtranscript pipeline with the following parameter: -mode multi [75]. Briefly, TEtranscript pipeline counts both uniquely and multi-mapped reads mapped to annotated transposons and genes to attribute transcript abundance. A given TE was considered transcribed only if reads mapped all along the element length, in order to avoid redundancy [75]. Levels of expression were calculated using Transcript per Million (TPM) normalization which corresponds to the normalization of read counts with transposon (or gene) length per million. We considered a TE as ‘expressed’ if TPM > 0.

### Sequence identification and comparison of DNA methyltransferases

All analyzed genomes and isolates used in this study are listed in Table S1 and originate from the following studies [32,33,77–81,34–37,48,49,71,76]. We identified homologs of *dim2* using the predicted ‘deRIPed’ protein sequence of *Z. tritici* IPO323 [25] as a template. To compare the sequence identity between active and inactive *dim2* copies in the genome, we used *dim2* of isolate Zt469 as a reference and performed pairwise comparisons. To identify additional putative DNA methyltransferases, we used the DNA methyltransferase domain of the Dim2 protein as query. BLAST searches, phylogenetic trees and alignments were performed using Geneious ‘Blast’, Geneious ‘Alignment’ and Geneious ‘Map to reference’ (Geneious version 10.2.4 (http://www.geneious.com, [82]). A distance tree based on the nucleotide sequence of *dim2, rid* and *dnmt5* was generated with the following settings: alignment type: global alignment with free end gaps, cost matrix: 65 % similarity, genetic distance: Jukes-Cantor, tree build method: neighbor joining, outgroup: *Zymoseptoria passerinii* (Zpa63).

### Generation of *dim2* deletion and integration strains

We transformed the *Z. tritici* isolates Zt09 and Zt10 using an *Agrobacterium tumefaciens*-mediated transformation (ATMT) protocol as previously described [83]. Briefly, we created plasmids containing the integration (pES189) or deletion (pES188) constructs using Gibson assembly [84] (S8 Table). Both constructs contained the hygromycin resistance cassette as selection marker. Plasmids were amplified in *E. coli* TOP10 cells and sequenced to confirm correct assembly of the constructs followed by electroporation of *A. tumefaciens* strain AGL1. *Z. tritici* strains were transformed with the respective *A. tumefaciens* strains by co-incubation for 3-4 days at 18°C on induction medium. Following co-incubation, the strains were grown on selection medium containing hygromycin to select for integration of the constructs and cefotaxime to eliminate *A. tumefaciens*. Single *Z. tritici* colonies were selected, streaked out twice and the correct integration of the construct was verified by PCR and by Southern blot analyses (S3 Fig).

### Bisulfite treatment and sequencing

We extracted high molecular weight DNA of three biological replicates of Zt10 and Zt09 and three independent transformants of Zt10Δ*dim2* and Zt09::*dim2* as described previously [70]. Genomic DNA was sent to the Max Planck Genome Centre in Cologne, Germany (https://mpgc.mpipz.mpg.de/home/) for bisulfite treatment and sequencing. Lambda DNA was used as spike-in (∼1%) to determine conversion efficiency. Genomic DNA was fragmented with a COVARIS S2 and an Illumina-compatible library was prepared with the NEXTflex Bisulfite Library Prep Kit for Illumina Sequencing (Bioo Scientific/PerkinElmer, Austin, Texas, U.S.A). Bisulfite conversion was performed using the EZ DNA Methylation Gold Kit (Zymo Research, Irvine, CA). Illumina sequencing was performed on a HiSeq3000 machine with paired-end 150-nt read mode (S9 Table).

To confirm bisulfite sequencing results, we treated genomic DNA of the fungal strains and, as a control for conversion efficiency, the Universal Methylated DNA Standard (Zymo Research, Irvine, CA) (∼100 ng) with bisulfite using the EZ DNA Methylation-Lightning Kit (Zymo Research, Irvine, CA) according to manufacturer’s instructions. Two representative loci per fungal strain and the human MLH1 for the control were amplified by PCR (ZymoTaq PreMix, Zymo Research, Irvine, CA) using specifically designed primers for amplification of the bisulfite-treated DNA (designed using the ‘Bisulfite Primer Seeker’ (Zymo Research, Irvine, CA), S8 Table). PCR products were cloned using the TOPO TA kit (Thermo Fisher Scientific) and Sanger sequenced (Eurofins Genomics, Ebersberg, Germany). Analysis of sequenced bisulfite-converted DNA sequences was performed using Geneious software version 10.2.4 [82].

### Data analysis of whole genome bisulfite sequencing data

A list of software and input commands used in our analyses is provided in the S1 text. Reads were quality filtered using Trimmomatic [73] and subsequently mapped using Bismark [85]. Duplicate reads were removed. Methylated sites were extracted using the bismark_methylation_extractor applying the –no_overlap and CX options. We considered sites methylated, if at least 4 reads and ≥ 50 % of reads supported methylation. We then extracted these sites (see text S1) in CG, CHG or CHH contexts for further analysis. Bedtools [86] was used to correlate methylation and genomics features.

### Southern blots to detect DNA methylation and confirm *dim2* mutant strains

To detect presence or absence of 5mC, we performed Southern blots according to standard protocols [87]. Genomic DNA was extracted using a standard phenol-chloroform extraction method [88]. The same amount of DNA and enzymes (*Bfu*CI, *Dpn*I, *Dpn*II; 25 units, New England Biolabs, Frankfurt, Germany) was used as input for the different restriction digests to make restriction patterns comparable between enzymes and *Z. tritici* isolates for the detection of DNA methylation. Probes were generated with the PCR DIG labeling Mix (Roche, Mannheim, Germany) following the manufacturer’s instructions and chemiluminescent signals were detected using the GelDocTM XR+ system (Bio-Rad, Munich, Germany).

### Phenotypic assay *in vitro*

Spores were diluted in water (107 cells/mL and tenfold dilution series to 1,000 cells/mL) and three µL of the spore suspension dilutions were pipetted on plates and incubated for ten days. To test for responses to different stress conditions *in vitro*, YMS plates containing NaCl (0.5 M and 1 M), sorbitol (1 M and 1.5 M), Congo Red (300 µg/mL and 500 µg/mL), H2O2 (1.5 mM and 2 mM), methyl methanesulfonate (0.01% and 0.005%) and two plates containing only YMS were prepared. All plates were incubated at 18°C, except for one of the YMS plates that was incubated at 28°C.

### Phenotypic assay on wheat

Seedlings of the wheat cultivar Obelisk (Wiersum Plantbreeding BV, Winschoten, The Netherlands) were pre-germinated on wet sterile Whatman paper for four days under normal growth conditions (16 h at light intensity of ∼200 µmol/m-2s-1 and 8 h darkness in growth chambers at 20°C with 90% humidity) followed by potting and further growth for additional seven days. Marked areas on the second leaves (30 leaves per strain) were inoculated with a spore suspension of 10^7^ cells/mL in H2O and 0.1 % Tween 20. Mock controls were treated with H2O and 0.1 % Tween 20 only. 23 days post inoculation treated leaves were analyzed for infection symptoms in form of necrosis and pycnidia. Evaluation was performed by assigning categories for necrosis and pycnidia coverage to each leaf (categories: 0 = 0%, 1 = 1-20%, 2 = 21-40%, 3 = 41-60%, 4 = 61-80%, 5 = 81-100%).

### Analysis of dinucleotide frequencies

TE sequences were extracted from the genome using bedtools getfasta [86]. k-mer frequencies in annotated TEs and masked genomes were determined using the software jellyfish [89]. Correspondence analyses were carried out and visualized in R [90] using the packages “FactoMineR” and “factoextra” [91].

### Mutation accumulation experiment and SNP analysis

A single colony derived directly from a plated dilution of frozen stock for Zt10 or IPO323 was resuspended in 1 mL YMS including 25% glycerol by 2 min vortexing on a VXR basic Vibrax at 2000 rpm, and 10–50 µL were re-plated onto a YMS agar plate. Forty replicates were produced. Cells were grown for 7 days at 18°C until a random colony (based on vicinity to a prefixed position on the plate) derived from a single cell was picked and transferred to a new plate as described above. The transfers were conducted for one year (52 times) before the DNA of a randomly chosen colony of each replicate was extracted and sequenced. Sequencing and library preparation were performed at the Max Planck Genome Center in Cologne, Germany (https://mpgc.mpipz.mpg.de/home/). Sequencing was performed on an Illumina HiSeq2500 machine obtaining paired-end 250-nt reads (S9 Table). Paired-end reads were quality filtered (Trimmomatic), mapped (bowtie2) and SNPs were called using samtools mpileup (see text S1).

## Data availability

Sequencing raw reads (FASTQ files) of all bisulfite and genomic data and the SMRT genome assemblies generated in this study are available online at Sequence Read Archive (SRA) under BioProject ID PRJNA614493. Previously published genome assemblies are available under BioProject PRJEB33986 [33], PRJNA638605, PRJNA639021 (Zpa63), PRJNA638553 (Zb87), PRJNA638515 (Zp13), PRJNA638382 (Za17), https://doi.org/10.5281/zenodo.3820378 [71], PRJNA414407 (Zt05 and Zt10) [36]. The genome sequence of the reference isolate IPO323 (Zt09) is available at: http://genome.jgi.doe.gov/Mycgr3/Mycgr3.home.html [26].

## Acknowledgements

Research in the lab of EHS is supported by the State of Schleswig-Holstein, the Max Planck Society and CIFAR. Research in the lab of MF is supported by NSF grant (MCB1818006). MM is supported by the German Research Foundation (DFG, MO 3755/1-1). We thank Bruce A. McDonald for providing Iranian *Z. tritici* isolates, Fatemeh Salimi for sampling Iranian *Z. tritici* isolates, and Kathrin Happ, Anja Lachner, and Maja Stralucke for assistance with experiments.

## Competing Interests

The authors declare no competing interests.

## Supplementary Information

### Supplementary Tables

**S1 Table**. List of *Z. tritici, Z. ardabiliae, Z. brevis* and *Z. passerinii* isolates used in this study. Listed are collection date, origin, information about the genome assembly and presence/absence of *dim2* in each isolate.

**S2 Table**. Detected copies and pairwise identity of full-length (>3,000 bp) *dim2* copies in *Z. tritici* genomes. The *dim2* gene of the Iranian isolate Zt469 was used as query for the blast search.

**S3 Table**. Dinucleotide frequencies in transposable elements (TEs) and TE masked genomes in *Z. tritici* isolates and the sister species *Z. brevis, Z. ardabiliae* and *Z. passerinii*.

**S4 Table**. Genes overlapping TEs and their functional annotation in Zt09.

**S5 Table**. SNPs detected in Zt10 and IPO323 after 52 weeks of experimental evolution compared to the reference strain at the start of the experiment.

**S6 Table**. Genome-wide occurrence of 5mC and SNPs in Zt10. Listed are GC and AT content, number of 5mC sites (pooled data from WGBS of all three replicates) and number of SNPs (pooled data from all 40 replicates) in 500 bp windows.

**S7 Table**. PacBio genome assembly metrics of Iranian isolates Zt289 and Zt469.

**S8 Table**. List of all oligos and plasmids used in this study.

**S9 Table**. Overview of Illumina sequencing data, detected 5mC sites, and SNPs.

**Supplementary text S1**. Software and commands used for bisulfite and genome data analysis to detect 5mC sites and SNPs.

### Supplementary Figures

**S1 Fig.**
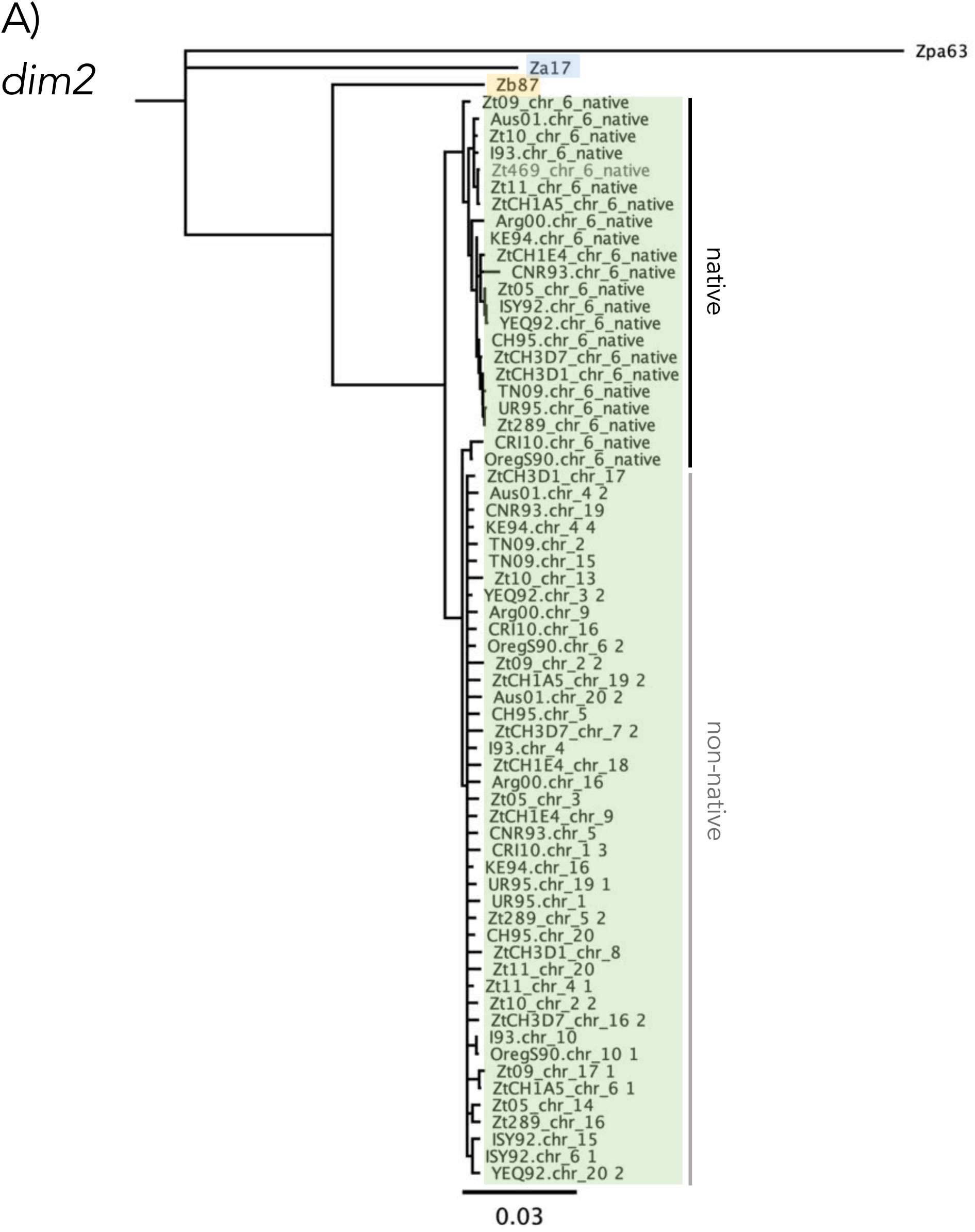
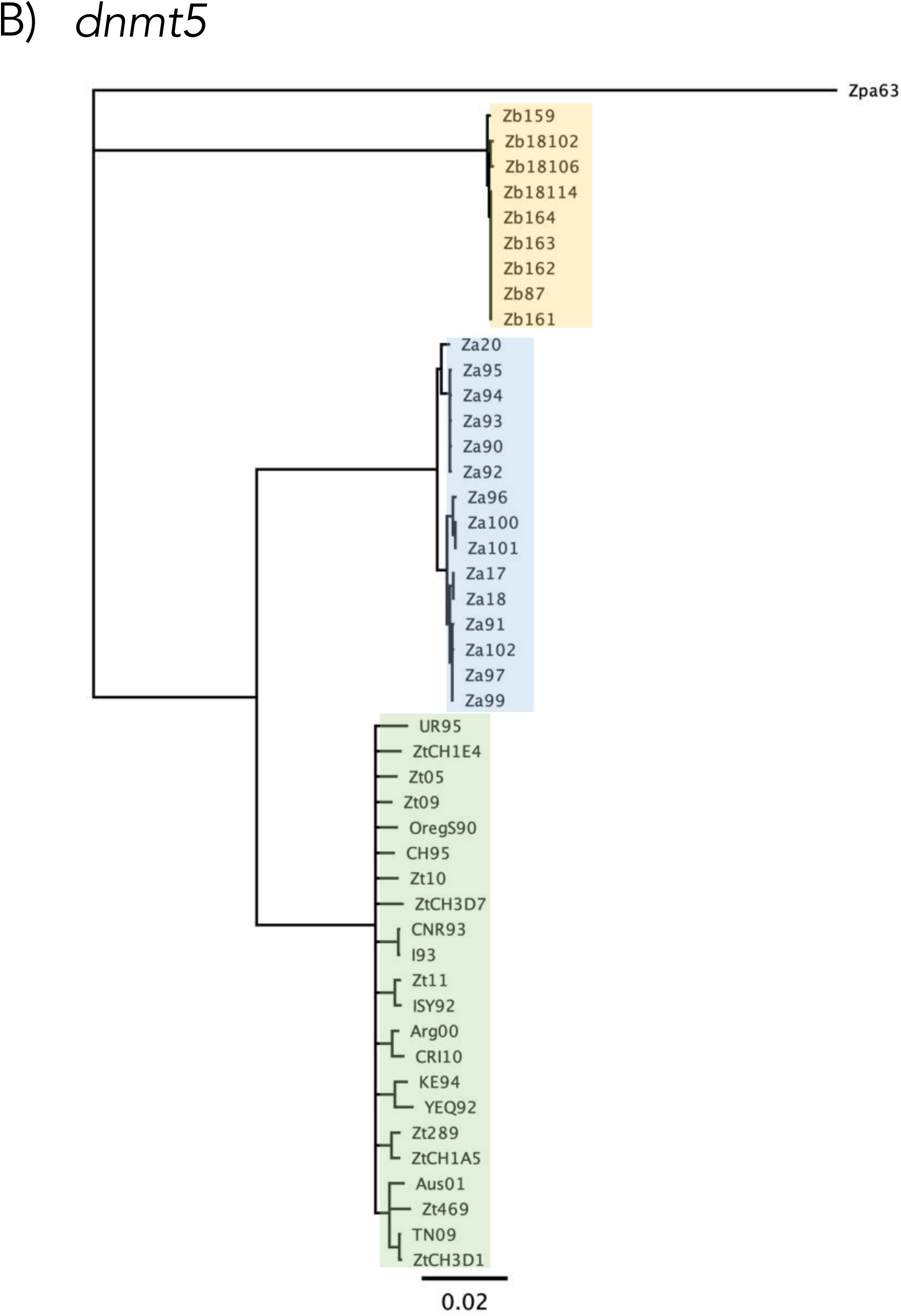
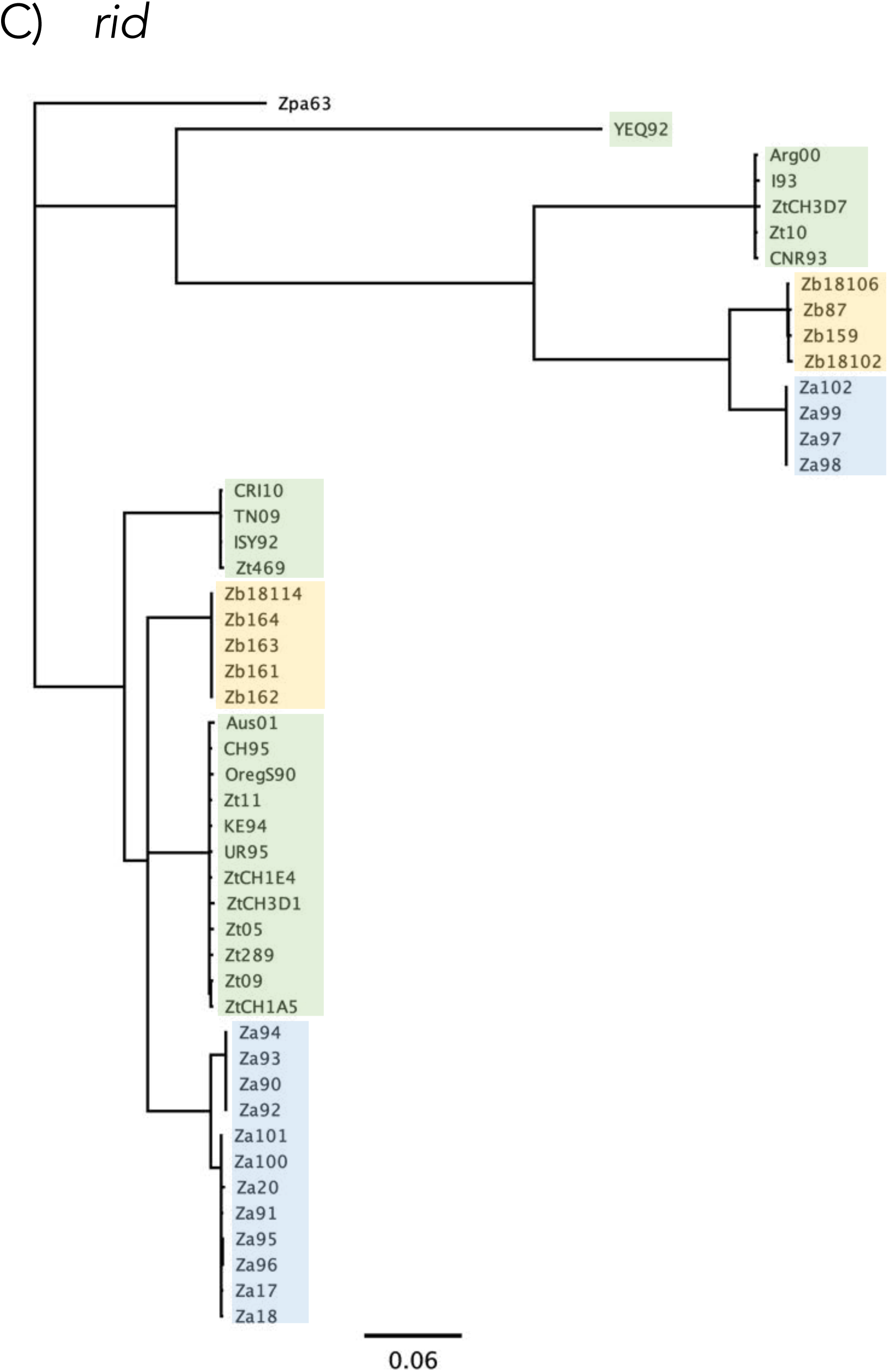
Phylogenetic trees of DNA sequences encoding the *dim2, dnmt5*, and *rid* genes. **A)** Phylogenetic tree based on alignments of “deRIPed” *dim2* alleles to the native, functional gene of Zt469. The *Z. tritici dim2* is distinct from the *Z. brevis* and *Z. ardabiliae* gene. The native *Z. tritici* copies, except for OregS90 and CRI10, and non-native copies form distinct clusters. Shown are two representative non-native copies per isolate. **B)** *Dnmt5* is present in all analyzed genomes and shows relatively little inter- or intraspecies diversity. **C)** The *rid gene* shows an exceptionally high inter- and intraspecies diversity with three highly distinct alleles present among genomes of *Z. tritici, Z. ardabiliae* and *Z. brevis*. Green background indicates *Z. tritici*, blue *Z. ardabiliae*, yellow *Z. brevis. Z. passerinii* is used as an outgroup.

**S2 Fig.**
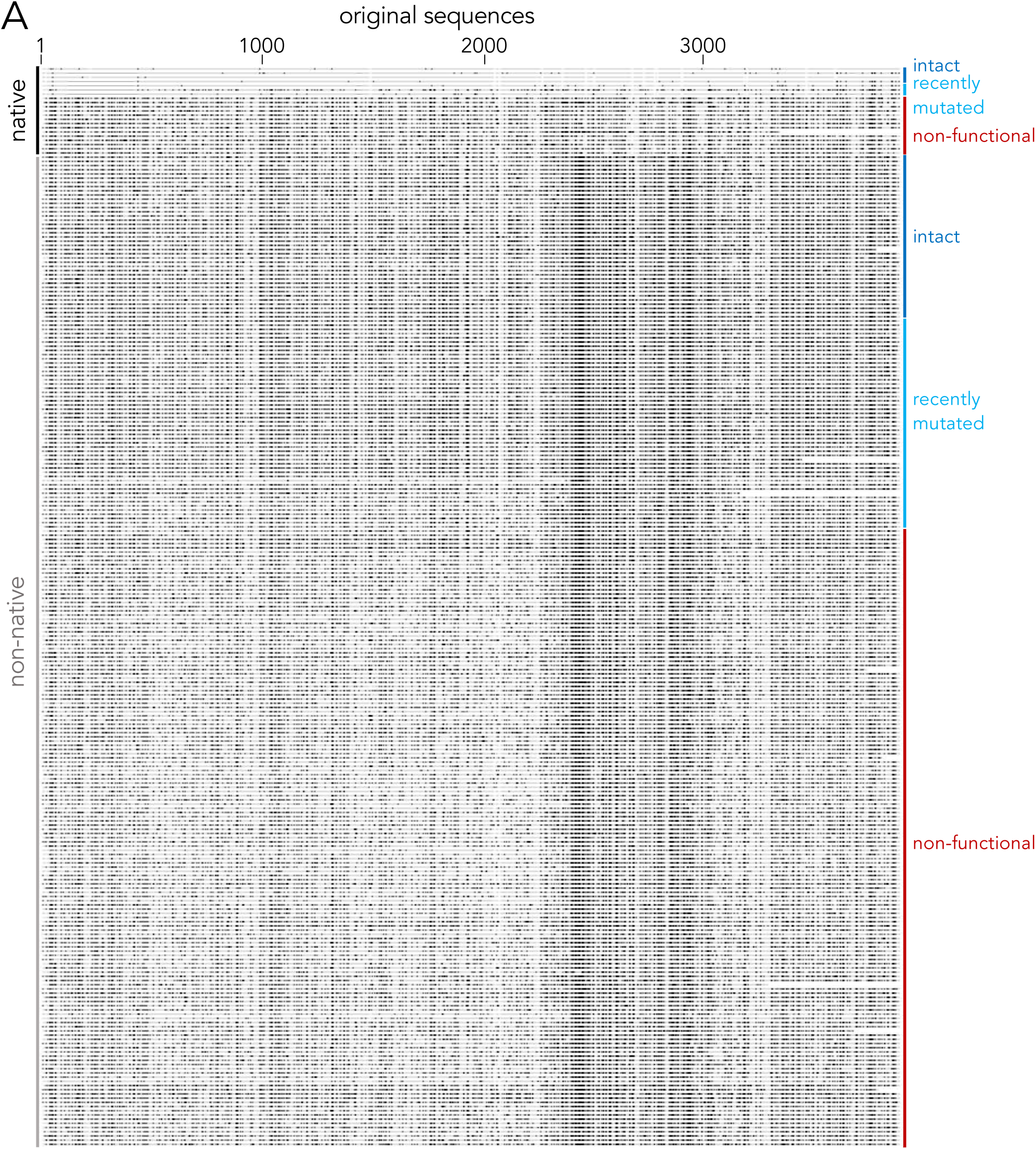
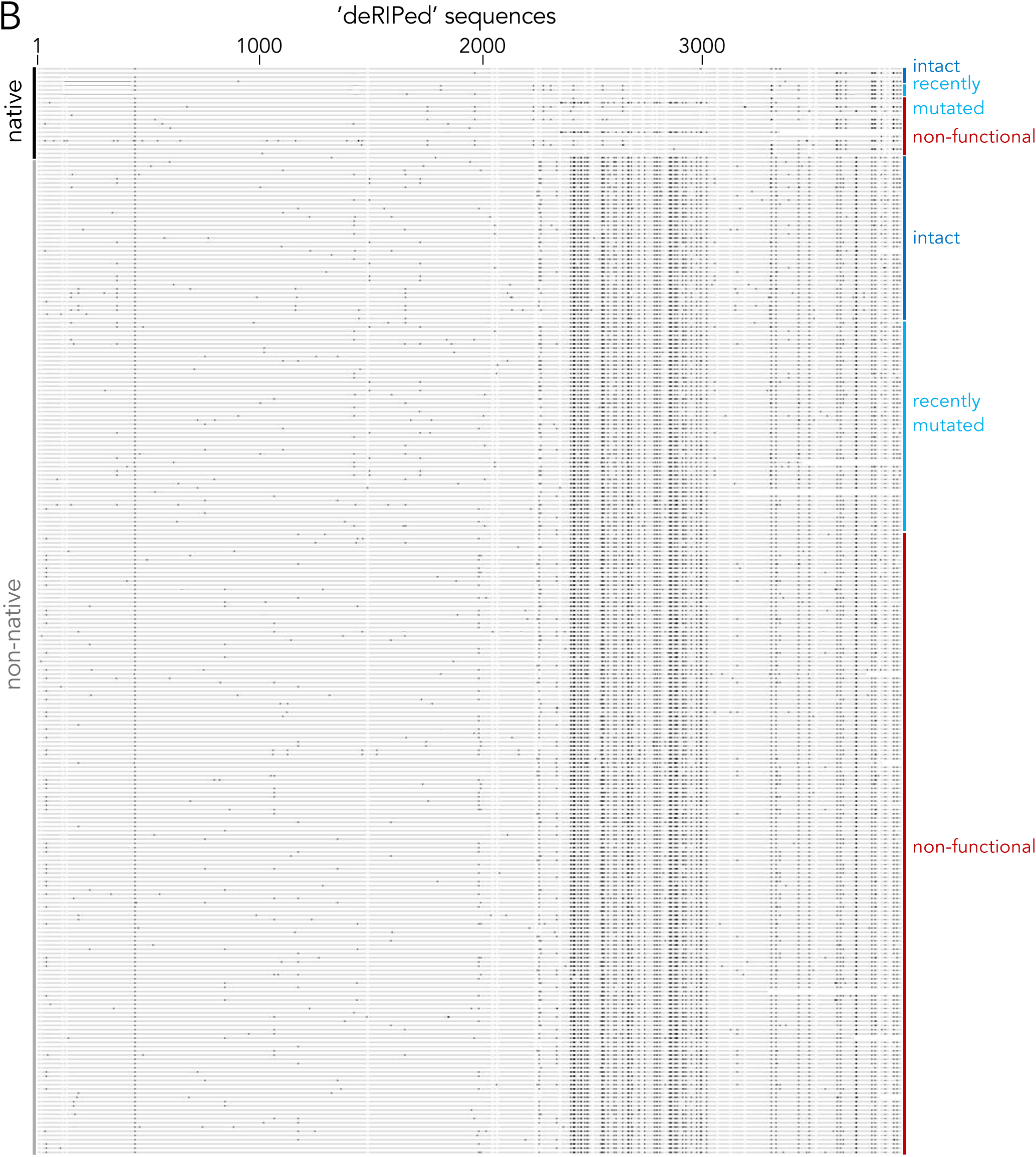
Alignment of original **(A)** and ‘deRIPed’ **(B)** full-length (> 3000 bp) *dim2* alleles compared to the functional allele of isolate Zt469. All native copies, (functional and non-functional, except for OregS90 and CRI10) lack mutations in the DNA methyltransferase domain (position ∼2,300–3,500) that are present in all non-native copies suggesting that the additional copies did not emerge from amplification of the native *dim2*. Black lines indicate differences to the reference.

**S3 Fig.**
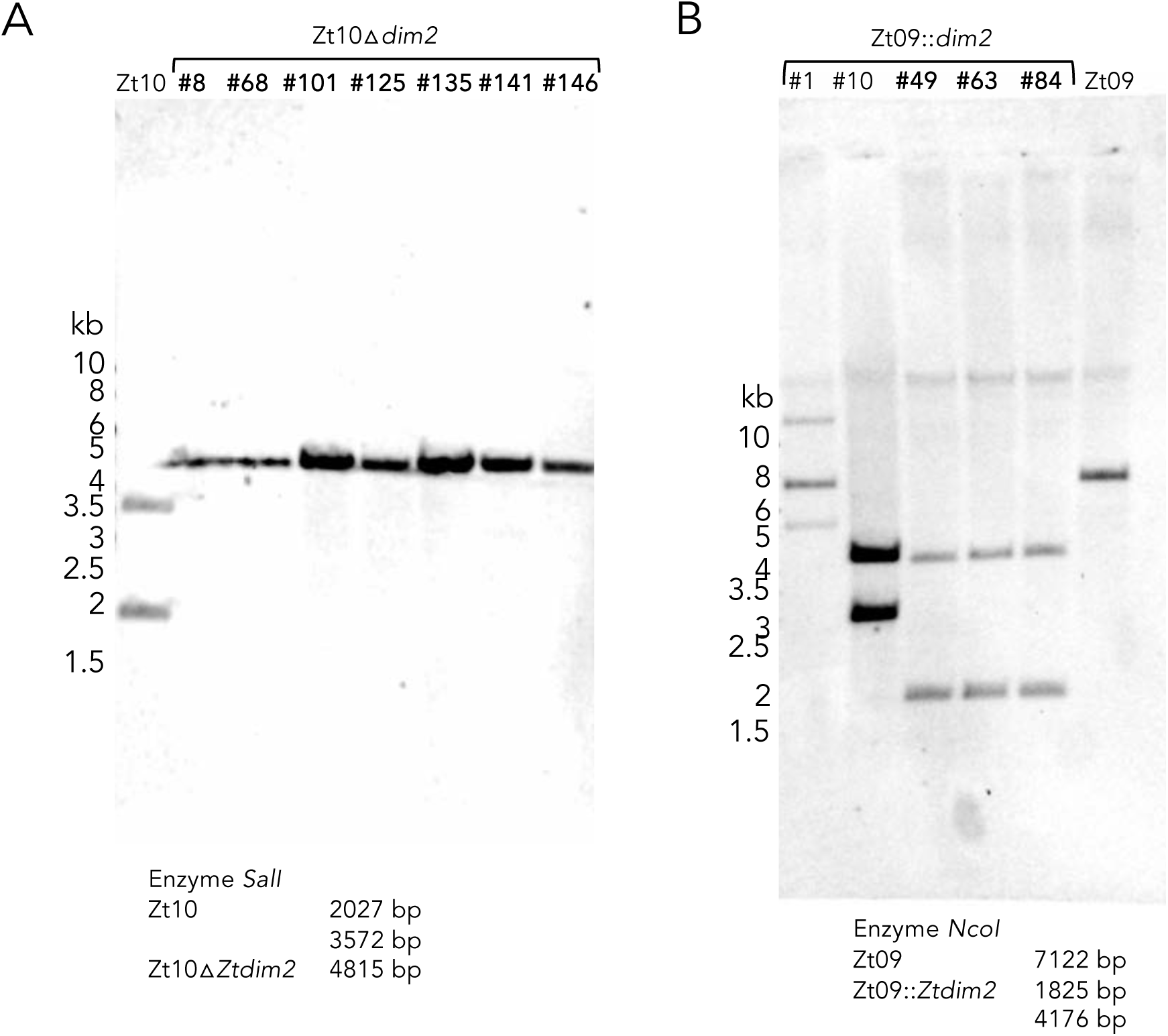
Southern blots to confirm the correct integration of **A)** the deletion construct for *dim2* in Zt10 (Zt10Δ*dim2*) and **B)** *dim2* (originating from Zt10) in Zt09 (Zt09::*dim2*). Three positive transformants (#49, #63 and #84) were found amongst the Zt09::*dim2* candidates, whereas all seven candidates for Zt10Δ*dim2* were verified (correct transformants are highlighted in bold).

**S4 Fig.**
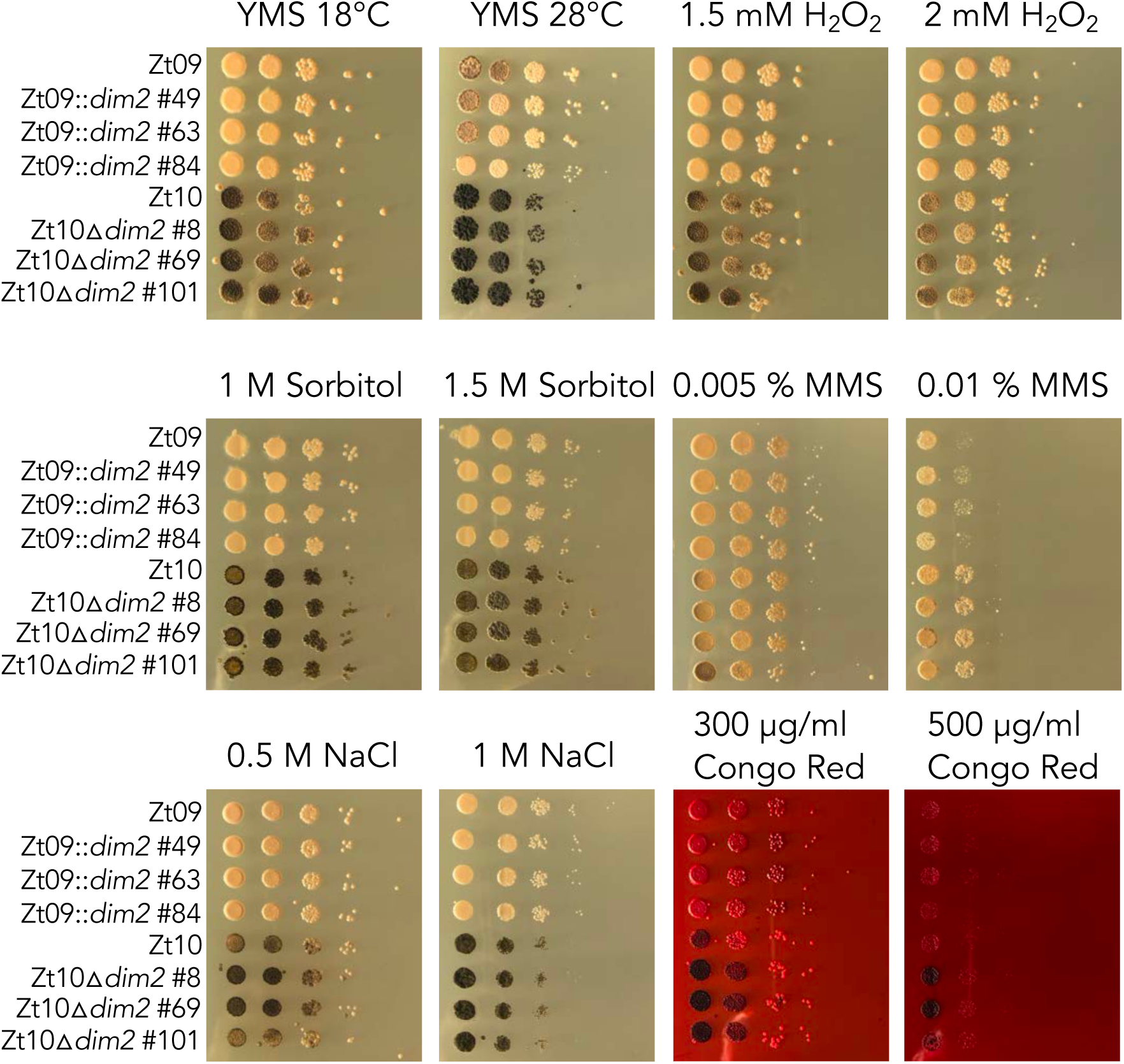
Phenotypic characterization of Zt10, Zt10Δ*dim2*, Zt09 and Zt09::*dim2 in vitro*. We compared growth phenotypes under different *in vitro* conditions including temperature, osmotic, oxidative, genotoxic and cell wall stress. We spotted spore dilutions of each reference isolate and three independent *dim2* mutant transformants on each plate. We did not detect any noticeable differences in growth between reference and mutant strains but differences between the different *Z. tritici* isolates Zt09 and Zt10 that were previously described [36].

**S5 Fig.**
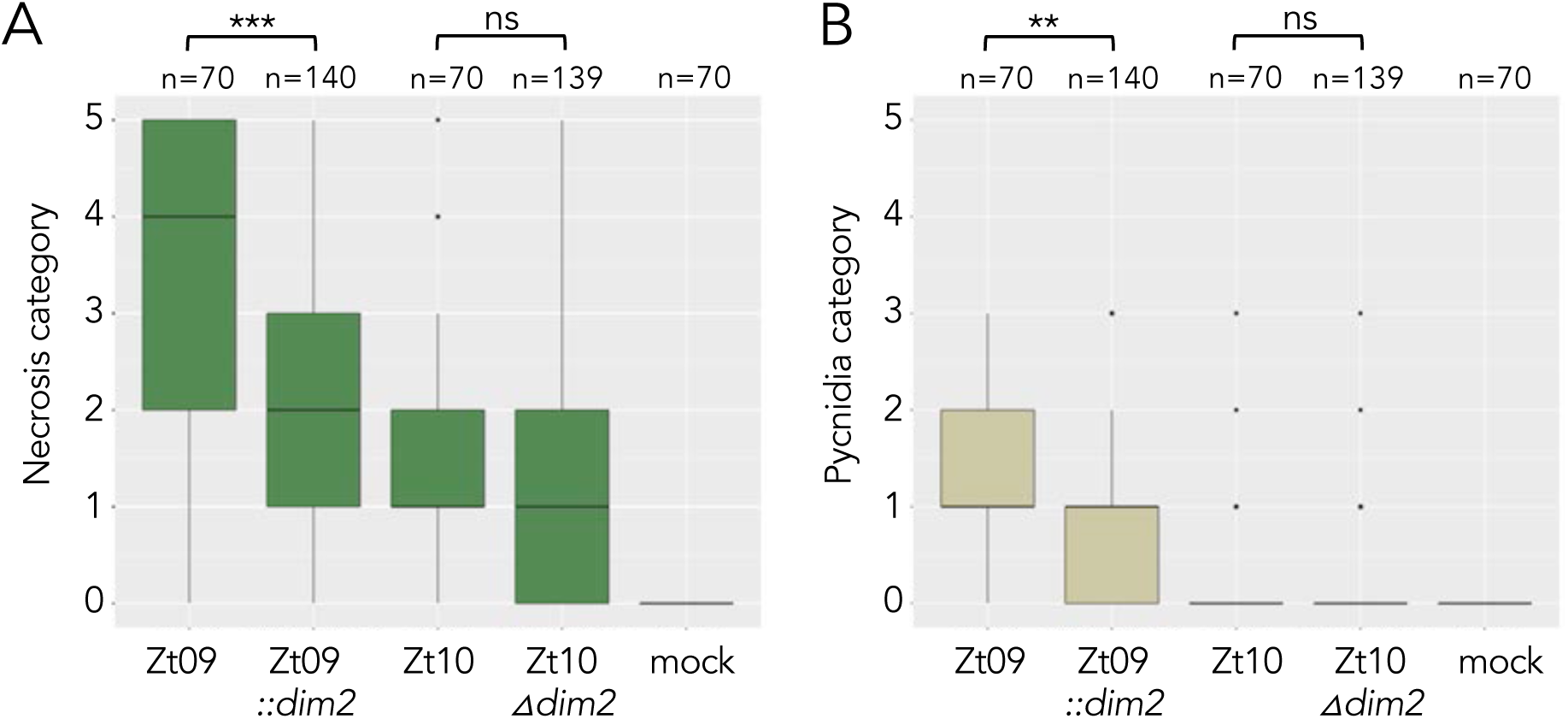
Results of wheat infection experiments with *Z. tritici* isolates Zt09, Zt10 and the mutants Zt09::*dim2* and Zt10Δ*dim2* on wheat. **A)** Zt09::*dim2* strains show significantly less necrotic lesions compared to Zt09 (*** Wilcoxon rank-sum test, *p*-value = 2.178 x 10^−7^) while there is no significant difference in the quantities of necrotic lesions caused by Zt10 and Zt10Δ*dim2* strains. **B)** Coverage with pycnidia is significantly reduced between Zt09 and Zt09::*dim2* (** *p*-value = 0.003301) but not between Zt10 and Zt10Δ*dim2* strains. Categories for necrotic lesion and pycnidia coverage: 0 = 0%, 1 = 1-20%, 2 = 21-40%, 3 = 41-60%, 4 = 61-80%, 5 = 81-100%.

**S6 Fig.**
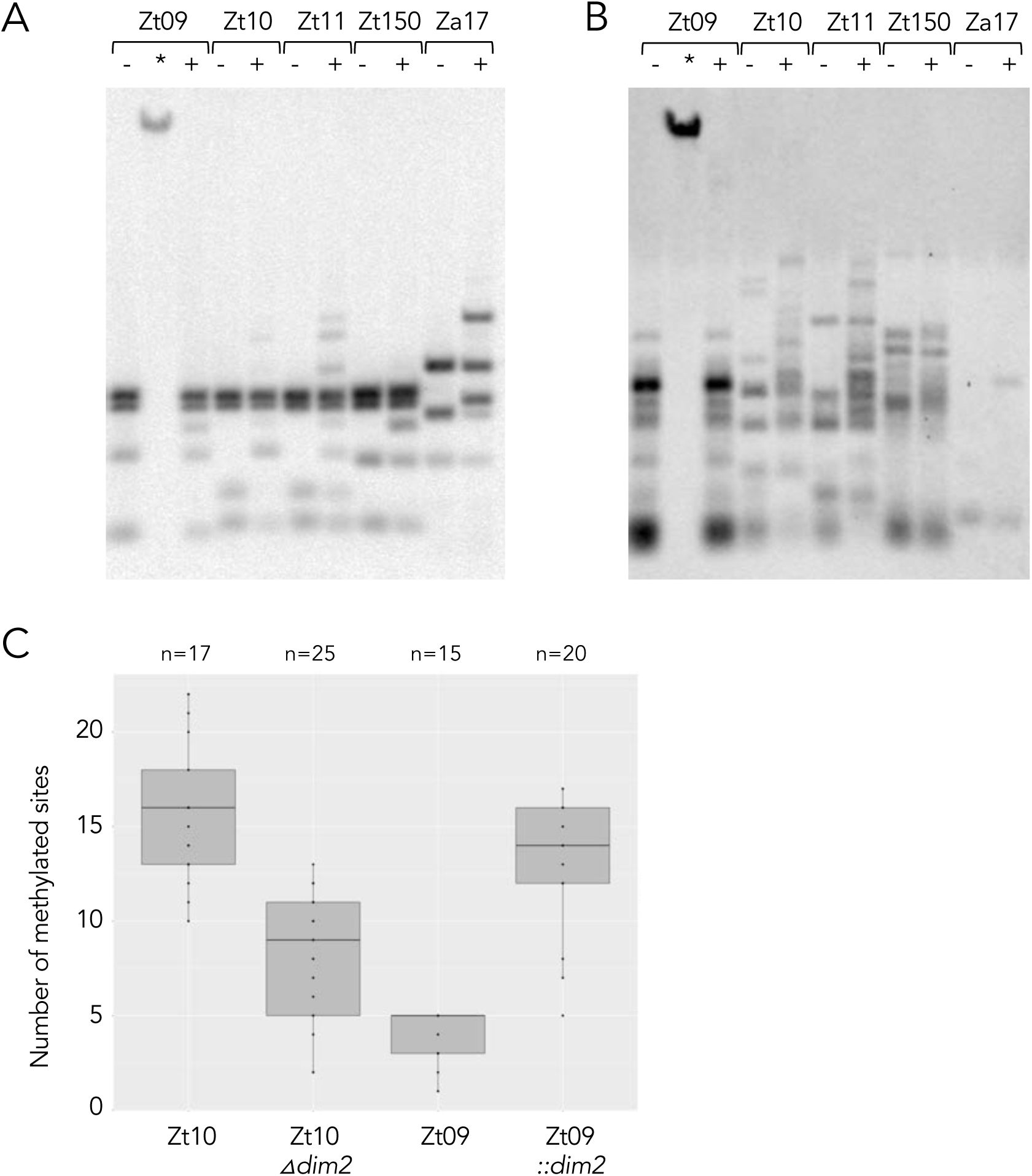
Detection of cytosine methylation by enzymatic digestion and Southern blot and confirmation of bisulfite sequencing results by PCR and Sanger sequencing. Restriction enzyme analysis followed by Southern blots using the rDNA spacer **(A)** or a retrotransposon RIL2 **(B)** as probe. In addition to the cytosine methylation sensitive (*Bfu*CI, **+**) or insensitive (*Dpn*II, **-**) enzymes, we used *Dpn*I (**\***) to test for the presence of adenine methylation. Zt09 and Zt150 contain a non-functional *dim2*, whereas the Iranian strains Zt10, Zt11 and the *Z. ardabiliae* strain Za17 have an intact copy. In all strains with a functional *dim2* we see a clear difference between the restrictions with *Bfu*CI and *Dpn*II, indicating the presence of DNA methylation. In Zt09 and Zt150 this difference is not detectable except for one band that is not restricted in both strains in the rDNA blot. The genomic DNA of Zt09 treated with *Dpn*I is not digested suggesting absence of 6mA methylation in these regions. **C)** Confirmation of bisulfite sequencing by an independent bisulfite treatment followed by PCR of target loci and Sanger sequencing. Two target loci (repeated region that showed 5mC signals in all isolates) were chosen per isolate, shown are the number of detected 5mC sites in these loci (size of loci ∼ 180 – 260 bp). N is the number of cloned PCR products that were sequenced. Based on these data we can confirm the presence of 5mC methylation in all isolates and a lower frequency in absence of *dim2*.

**S7 Fig.**
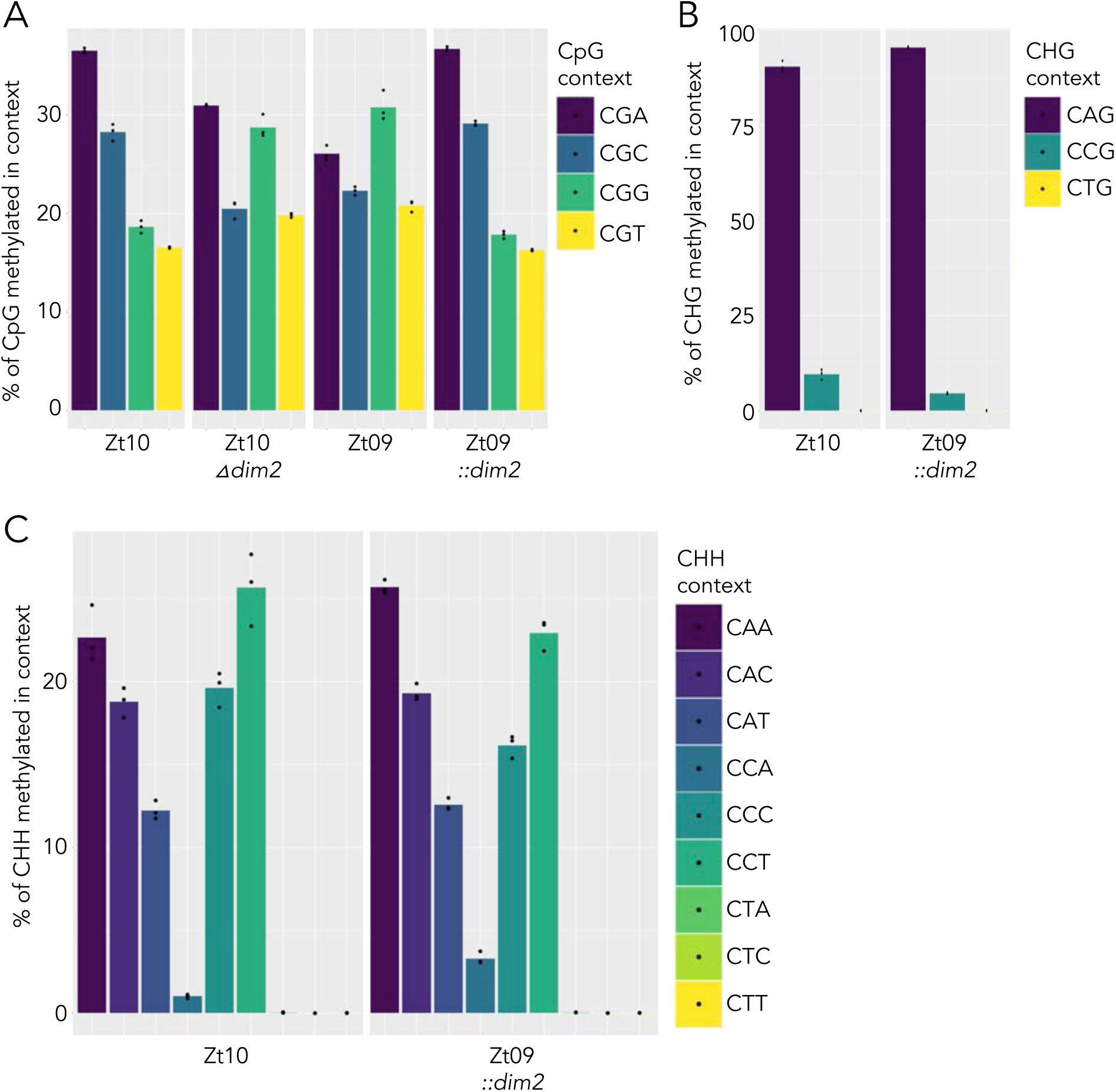
5mC site preferences in CG, CHG and CHH sequence contexts. **A)** CGA and CGC sites are preferred in CG contexts in presence of *dim2*, while CGA and CGG are slightly preferred targets in absence of *dim2*. **B)** Among CHG sites, CAG sites are the predominant target sites of 5mC. For CHG and CHH contexts only data for Zt10 and Zt09::*dim2* are shown, as there is no detectable methylation outside of CG contexts in absence of *dim2*. **C)** In CHH contexts, CA sites followed by CC sites have the highest methylation frequency. CT sites are almost completely devoid of 5mC.

**S8 Fig.**
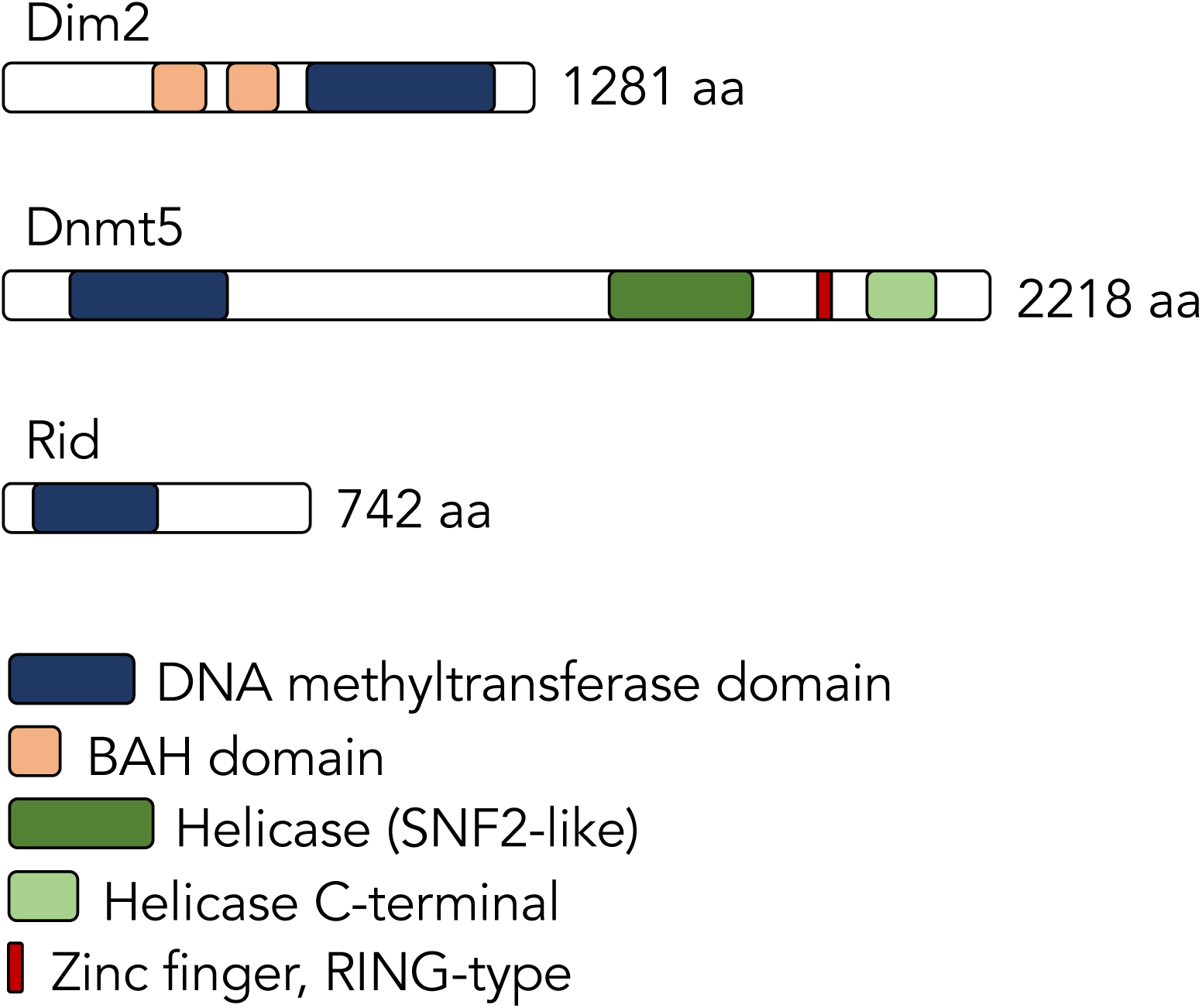
Predicted domains of the three putative DNA methyltransferase proteins Dim2, Dnmt5 and Rid identified in the *Zymoseptoria* genomes. Protein sequences were analyzed with SMART [92], NCBI Blast and InterProScan [93].

## Notes

### Competing Interest Statement

The authors have declared no competing interest.

## References

1. Jones PA. Functions of DNA methylation: Islands, start sites, gene bodies and beyond. Nat Rev Genet. 2012;13: 484–492. doi:10.1038/nrg3230

2. Slotkin RK, Martienssen R. Transposable elements and the epigenetic regulation of the genome. Nat Rev Genet. 2007;8: 272–85. doi:10.1038/nrg2072

3. Martienssen RA, Colot V. DNA Methylation and Epigenetic Inheritance in Plants and Filamentous Fungi. Science. 2001;293: 1070–1074. doi:10.1126/science.293.5532.1070

4. Zemach A, McDaniel IE, Silva P, Zilberman D. Genome-Wide Evolutionary Analysis of Eukaryotic DNA Methylation. Science. 2010;238: 916–919.

5. Lyko F. The DNA methyltransferase family: A versatile toolkit for epigenetic regulation. Nat Rev Genet. 2018;19: 81–92. doi:10.1038/nrg.2017.80

6. Robert M, Morin S, Beaulieu N, Gauthier F, Chute IC, Barsalou A, et al. DNMT1 is required to maintain CpG methylation and aberrant gene silencing in human cancer cells. Nat Genet. 2003;33: 61–65. doi:10.1038/ng1068

7. Gruenbaum Y, Cedar H, Kazin A. Substrate and sequence specificity of a eukaryotic DNA methylase. Nature. 1982;295: 620–622.

8. Leonhardt H, Page AW. A Targeting Sequence Directs DNA Methyltransferase to Sites of DNA Replication in Mammalian. Cell. 1992;71: 865–873.

9. Okano M, Bell DW, Haber DA, Li E. DNA Methyltransferases Dnmt3a and Dnmt3b Are Essential for De Novo Methylation and Mammalian Development. Cell. 1999;99: 247–257.

10. Cao X, Jacobsen SE. Role of the Arabidopsis DRM Methyltransferases in De Novo DNA Methylation and Gene Silencing. Curr Biol. 2002;12: 1138–1144. doi:https://doi.org/10.1016/S0960-9822(02)00925-9

11. Cao X, Springer NM, Muszynski MG, Phillips RL, Kaeppler S, Jacobsen SE. Conserved plant genes with similarity to mammalian de novo DNA methyltransferases. Proc Natl Acad Sci. 2000;97: 4979–4984.

12. Jeltsch A, Jurkowska RZ. New concepts in DNA methylation. Trends Biochem Sci. 2014;39: 310–318. doi:10.1016/j.tibs.2014.05.002

13. Goyal R, Reinhardt R, Jeltsch A. Accuracy of DNA methylation pattern preservation by the Dnmt1 methyltransferase. Nucleic Acids Res. 2006;34: 1182–1188. doi:10.1093/nar/gkl002

14. Zolan ME, Pukkila PJ. Inheritance of DNA methylation in Coprinus cinereus. Mol Cell Biol. 1986;6: 195–200. doi:10.1128/mcb.6.1.195

15. Kouzminova E, Selker EU. Dim-2 encodes a DNA methyltransferase responsible for all known cytosine methylation in Neurospora. EMBO J. 2001;20: 4309–4323. doi:10.1093/emboj/20.15.4309

16. Goll MG, Bestor TH. Eukaryotic Cytosine Methyltransferases. Annu Rev Biochem. 2005;74: 481–514. doi:10.1146/annurev.biochem.74.010904.153721

17. Freitag M, Williams RL, Kothe GO, Selker EU. A cytosine methyltransferase homologue is essential for repeat-induced point mutation in Neurospora crassa. Proc Natl Acad Sci. 2002;99: 8802–8807. doi:10.1073/pnas.132212899

18. Malagnac F, Wendel B, Goyon C, Faugeron G, Zickler D, Rossignol JL, et al. A gene essential for de novo methylation and development in ascobolus reveals a novel type of eukaryotic DNA methyltransferase structure. Cell. 1997;91: 281–290. doi:10.1016/S0092-8674(00)80410-9

19. Gladyshev E, Kleckner N. DNA sequence homology induces cytosine-to-thymine mutation by a heterochromatin-related pathway in Neurospora. Nature Genetics. 2017;49: 887–894. doi:10.1038/ng.3857

20. Huff JT, Zilberman D. Dnmt1-independent CG methylation contributes to nucleosome positioning in diverse eukaryotes. Cell. 2014;156: 1286–1297. doi:10.1016/j.cell.2014.01.029

21. Catania S, Dumesic PA, Pimentel H, Nasif A, Stoddard CI, Burke JE, et al. Evolutionary Persistence of DNA Methylation for Millions of Years after Ancient Loss of a De Novo Methyltransferase. Cell. 2020;180: 263-277.e20. doi:10.1016/j.cell.2019.12.012

22. Bewick AJ, Hofmeister BT, Powers RA, Mondo SJ, Grigoriev IV, James TY, et al. Diversity of cytosine methylation across the fungal tree of life. Nat Ecol Evol. 2019; doi:10.1038/s41559-019-0810-9

23. Ruesch CE, Ramakrishnan M, Park J, Li N, Chong HS, Zaman R, et al. The Histone H3 Lysine 9 Methyltransferase DIM-5 Modifies Chromatin at frequency and Represses Light-Activated Gene Expression. G3: GENES, GENOMES, GENETICS. 2015;5: 93–101. doi:10.1534/g3.114.015446

24. Xue Z, Ye Q, Anson SR, Yang J, Xiao G, Kowbel D, et al. Transcriptional interference by antisense RNA is required for circadian clock function. Nature. 2014/08/17. 2014;514: 650–653. doi:10.1038/nature13671

25. Dhillon B, Cavaletto JR, Wood KV, Goodwin SB. Accidental amplification and inactivation of a methyltransferase gene eliminates cytosine methylation in Mycosphaerella graminicola. Genetics. 2010;186: 67–77. doi:10.1534/genetics.110.117408

26. Goodwin SB, M’barek SB, Dhillon B, Wittenberg AHJ, Crane CF, Hane JK, et al. Finished genome of the fungal wheat pathogen Mycosphaerella graminicola reveals dispensome structure, chromosome plasticity, and stealth pathogenesis. PLOS Genet. 2011;7: e1002070. doi:10.1371/journal.pgen.1002070

27. Linde CC, Zhan J, McDonald BA. Population Structure of Mycosphaerella graminicola: From Lesions to Continents. Phytopathology. 2002;92: 946–955. doi:10.1094/PHYTO.2002.92.9.946

28. Zhan J, Pettway RE, McDonald BA. The global genetic structure of the wheat pathogen Mycosphaerella graminicola is characterized by high nuclear diversity, low mitochondrial diversity, regular recombination, and gene flow. Fungal Genet Biol. 2003;38: 286–297. doi:10.1016/S1087-1845(02)00538-8

29. Hartmann FE, Sánchez-Vallet A, McDonald BA, Croll D. A fungal wheat pathogen evolved host specialization by extensive chromosomal rearrangements. ISME J. 2017;11: 1189–1204. doi:10.1038/ismej.2016.196

30. Oggenfuss U, Badet T, Wicker T, Hartmann FE, Singh NK, Abraham LN, et al. A population-level invasion by transposable elements in a fungal pathogen. bioRxiv. 2020; 2020.02.11.944652. doi:10.1101/2020.02.11.944652

31. Lorrain C, Feurtey A, Möller M, Haueisen J, Stukenbrock E. Dynamics of transposable elements in recently diverged fungal pathogens: lineage-specific transposable element content and efficiency of genome defences. bioRxiv. 2020; 2020.05.13.092635. doi:10.1101/2020.05.13.092635

32. Feurtey A, Stevens DM, Stephan W, Stukenbrock EH. Interspecific gene exchange introduces high genetic variability in crop pathogen. Genome Biol Evol. 2019;evz224. doi:10.1093/gbe/evz224

33. Badet T, Oggenfuss U, Abraham L, McDonald BA, Croll D. A 19-isolate reference-quality global pangenome for the fungal wheat pathogen Zymoseptoria tritici. BMC Biol. 2020;18: 12. doi:10.1186/s12915-020-0744-3

34. Plissonneau C, Stürchler A. The Evolution of Orphan Regions in Genomes of a Fungal Pathogen of Wheat. mBio. 2016;7: e01231–16. doi:10.1128/mBio.01231-16.

35. Plissonneau C, Hartmann FE, Croll D. Pangenome analyses of the wheat pathogen Zymoseptoria tritici reveal the structural basis of a highly plastic eukaryotic genome. BMC Biol. 2018;16: 5. doi:10.1186/s12915-017-0457-4

36. Haueisen J, Möller M, Eschenbrenner CJ, Grandaubert J, Seybold H, Adamiak H, et al. Highly flexible infection programs in a specialized wheat pathogen. Ecol Evol. 2019;9: 275–294. doi:10.1002/ece3.4724

37. Stukenbrock EH, Bataillon T, Dutheil JY, Hansen TT, Li R, Zala M, et al. The making of a new pathogen: Insights from comparative population genomics of the domesticated wheat pathogen Mycosphaerella graminicola and its wild sister species. Genome Res. 2011;21: 2157–2166. doi:10.1101/gr.118851.110

38. Galagan JE, Selker EU. RIP: The evolutionary cost of genome defense. Trends Genet. 2004;20: 417–423. doi:10.1016/j.tig.2004.07.007

39. Selker EU. PREMEIOTIC INSTABILITY OF REPEATED SEQUENCES IN NEUROSPORA CRASSA. Annu Rev Genet. 1990;24: 579–613. doi:10.1146/annurev.ge.24.120190.003051

40. Möller M, Schotanus K, Soyer JL, Haueisen J, Happ K, Stralucke M, et al. Destabilization of chromosome structure by histone H3 lysine 27 methylation. PLOS Genet. 2019;15: e1008093. doi.org/10.1371/journal.pgen.1008093

41. Tamaru H, Selker EU. A histone H3 methyltransferase controls DNA methylation in Neurospora crassa. Nature. 2001;414: 277–83. doi:10.1038/35104508

42. Jackson JP, Lindroth AM, Cao X, Jacobsen SE. Control of CpNpG DNA methylation by the KRYPTONITE histone H3 methyltransferase. Nature. 2002;416: 556–560. doi:10.1038/nature731

43. Du J, Johnson LM, Jacobsen SE, Patel DJ. DNA methylation pathways and their crosstalk with histone methylation. Nat Rev Mol Cell Biol. 2015;16. doi:10.1038/nrm4043

44. Dumesic PA, Homer CM, Moresco JJ, Pack LR, Shanle EK, Coyle SM, et al. Product Binding Enforces the Genomic Specificity of a Yeast Polycomb Repressive Complex. Cell. 2015;160: 204–218. doi:10.1016/j.cell.2014.11.039

45. Grandaubert J, Bhattacharyya A, Stukenbrock EH. RNA-seq Based Gene Annotation and Comparative Genomics of Four Fungal Grass Pathogens in the Genus Zymoseptoria Identify Novel Orphan Genes and Species-Specific Invasions of Transposable Elements. G3: GENES, GENOMES, GENETICS. 2015;5: g3.115.017731-. doi:10.1534/g3.115.017731

46. Selker EU, Stevens JN. DNA methylation at asymmetric sites is associated with numerous transition mutations. Proc Natl Acad Sci. 1985;82: 8114–8. doi:10.1073/pnas.82.23.8114

47. Holliday R, Grigg GW. DNA methylation and mutation. Mutat Res. 1993;285: 61–67.

48. Stukenbrock EH, Quaedvlieg W, Javan-Nikhah M, Zala M, Crous PW, McDonald BA. Zymoseptoria ardabiliae and Z. pseudotritici, two progenitor species of the septoria tritici leaf blotch fungus Z. tritici (synonym: Mycosphaerella graminicola). Mycologia. 2012;104: 1397–407. doi:10.3852/11-374

49. Quaedvlieg W, Kema GHJ, Groenewald JZ, Verkley GJM, Seifbarghi S, Razavi M, et al. Zymoseptoria gen. nov.: A new genus to accommodate Septoria-like species occurring on graminicolous hosts. Persoonia Mol Phylogeny Evol Fungi. 2011;26: 57–69. doi:10.3767/003158511X571841

50. McDonald MC, McGinness L, Hane JK, Williams AH, Milgate A, Solomon PS. Utilizing Gene Tree Variation to Identify Candidate Effector Genes in Zymoseptoria tritici. G3: GENES, GENOMES, GENETICS. 2016;6: 779–91. doi:10.1534/g3.115.025197

51. Montanini B, Chen P-Y, Morselli M, Jaroszewicz A, Lopez D, Martin F, et al. Non-exhaustive DNA methylation-mediated transposon silencing in the black truffle genome, a complex fungal genome with massive repeat element content. Genome Biol. 2014;15: 411. doi:10.1186/s13059-014-0411-5

52. Nakayashiki H, Ikeda K, Hashimoto Y, Tosa Y, Mayama S. Methylation is not the main force repressing the retrotransposon MAGGY in Magnaporthe grisea. Nucleic Acids Res. 2001;29: 1278–84. doi:10.1093/nar/29.6.1278

53. Duncan BK, Miller JH. Mutagenic deamination of cytosine residues in DNA. Nature. 1980;287: 560–561. doi:10.1038/287560a0

54. Kiefer C, Willing E, Jiao W, Sun H, Piednoël M, Hümann U, et al. Interspecies association mapping links reduced CG to TG substitution rates to the loss of gene body methylation. Nat Plants. 2019;5. doi:10.1038/s41477-019-0486-9

55. Mazur AK, Gladyshev E. Partition of Repeat-Induced Point Mutations Reveals Structural Aspects of Homologous DNA-DNA Pairing. Biophys J. 2018;115: 605–615. doi:10.1016/j.bpj.2018.06.030

56. Yang AS, Shen JC, Zingg JM, Mi S, Jones PA. HhaI and HpaII DNA methyltransferases bind DNA mismatches, methylate uracil and block DNA repair. Nucleic Acids Res. 1995;23: 1380–1387. doi:10.1093/nar/23.8.1380

57. Shen JC, Zingg JM, Yang AS, Schmutte C, Jones PA. A mutant HpaII methyltransferase functions as a mutator enzyme. Nucleic Acids Res. 1995;23: 4275–4282. doi:10.1093/nar/23.21.4275

58. Rosa AL, Folco HD, Mautino MR. In vivo levels of S-adenosylmethionine modulate C:G to T:A mutations associated with repeat-induced point mutation in Neurospora crassa. Mutat Res Mol Mech Mutagen. 2004;548: 85–95. doi:https://doi.org/10.1016/j.mrfmmm.2004.01.001

59. Mautino MR, Rosa AL. Analysis of Models Involving Enzymatic Activities for the Occurrence of C→T Transition Mutations During Repeat-Induced Point Mutation (RIP) in Neurospora crassa. J Theor Biol. 1998;192: 61–71. doi:https://doi.org/10.1006/jtbi.1997.0608

60. Chen L, MacMillan AM, Verdine GL. Mutational separation of DNA binding from catalysis in a DNA cytosine methyltransferase. J Am Chem Soc. 1993;115: 5318–5319. doi:10.1021/ja00065a063

61. Shen J-C, Rideout III WM, Jones PA. High frequency mutagenesis by a DNA methyltransferase. Cell. 1992;71: 1073–1080. doi:10.1016/S0092-8674(05)80057-1

62. Wang K-Y, James Shen C-K. DNA methyltransferase Dnmt1 and mismatch repair. Oncogene. 2004;23: 7898–7902. doi:10.1038/sj.onc.1208111

63. Freitag M. Histone Methylation by SET Domain Proteins in Fungi. Annu Rev Microbiol. 2017;36: 413–439. doi:10.1146/annurev-micro-102215-095757

64. Ikeda KI, Van Vu B, Kadotani N, Tanaka M, Murata T, Shiina K, et al. Is the fungus Magnaporthe losing DNA methylation? Genetics. 2013;195: 845–855. doi:10.1534/genetics.113.155978

65. Yang K, Liang L, Ran F, Liu Y, Li Z, Lan H, et al. The DmtA methyltransferase contributes to Aspergillus flavus conidiation, sclerotial production, aflatoxin biosynthesis and virulence. Sci Rep. 2016;6: 1–13. doi:10.1038/srep23259

66. Selmecki A, Forche A, Berman J. Genomic plasticity of the human fungal pathogen Candida albicans. Eukaryot Cell. 2010;9: 991–1008. doi:10.1128/EC.00060-10

67. Hu G, Wang J, Choi J, Jung WH, Liu I, Litvintseva AP, et al. Variation in chromosome copy number influences the virulence of Cryptococcus neoformans and occurs in isolates from AIDS patients. BMC Genomics. 2011;12: 526. doi:10.1186/1471-2164-12-526

68. Möller M, Stukenbrock EH. Evolution and genome architecture in fungal plant pathogens. Nat Rev Microbiol. 2017;15: 756. doi.org/10.1038/nrmicro.2017.76

69. Raffaele S, Kamoun S. Genome evolution in filamentous plant pathogens: why bigger can be better. Nat Rev Microbiol. 2012;10: 417–430. doi:10.1038/nrmicro2790

70. Allen GC, Flores-Vergara MA, Krasynanski S, Kumar S, Thompson WF. A modified protocol for rapid DNA isolation from plant tissues using cetyltrimethylammonium bromide. Nat Protoc. 2006;1: 2320–2325. doi:10.1038/nprot.2006.384

71. Feurtey A, Lorrain C, Croll D, Eschenbrenner C, Freitag M, Habig M, et al. Genome compartmentalization predates species divergence in the plant pathogen genus Zymoseptoria. BMC Genomics. 2020;21: 588. doi:10.1186/s12864-020-06871-w

72. Flutre T, Duprat E, Feuillet C, Quesneville H. Considering Transposable Element Diversification in De Novo Annotation Approaches. PLoS One. 2011;6: e16526. doi:10.1371/journal.pone.0016526

73. Bolger AM, Lohse M, Usadel B. Trimmomatic: A flexible trimmer for Illumina sequence data. Bioinformatics. 2014;30: 2114–2120. doi:10.1093/bioinformatics/btu170

74. Kim D, Langmead B, Salzberg SL. HISAT: a fast spliced aligner with low memory requirements. Nat Methods. 2015;12: 357–60. doi:10.1038/nmeth.3317

75. Jin Y, Hammell M. Analysis of RNA-Seq Data Using TE transcripts. Methods Mol Biol. 2018;1751: 153–167. doi:10.1007/978-1-4939-7710-9_11

76. Stukenbrock EH, Dutheil JY. Fine-scale recombination maps of fungal plant pathogens reveal dynamic recombination landscapes and intragenic hotspots. Genetics. 2018;208: 1209–1229. doi:10.1534/genetics.117.300502

77. Möller M, Habig M, Freitag M, Stukenbrock EH. Extraordinary Genome Instability and Widespread Chromosome Rearrangements During Vegetative Growth. Genetics. 2018;210 no. 2: 517–529. doi:10.1534/genetics.118.301050

78. Grandaubert J, Dutheil JY, Stukenbrock EH. The genomic determinants of adaptive evolution in a fungal pathogen. Evolution Letters. 2019; 299–312. doi:10.1002/evl3.117

79. Jürgens T, Linde CC, Mcdonald BA. Genetic structure of Mycosphaerella graminicola populations from Iran, Argentina and Australia. Eur J Plant Pathol. 2006; 115:223–23. doi:10.1007/s10658-006-9000-0

80. Daboussi MJ. Fungal transposable elements and genome evolution. Genetica. 1997;100: 253–260. doi:10.1023/A:1018354200997

81. Zhan J, Linde CC, Jürgens T, Merz U, Steinebrunner F, McDonald BA. Variation for neutral markers is correlated with variation for quantitative traits in the plant pathogenic fungus Mycosphaerella graminicola. Mol Ecol. 2005;14: 2683–2693. doi:10.1111/j.1365-294X.2005.02638.x

82. Kearse M, Moir R, Wilson A, Stones-Havas S, Cheung M, Sturrock S, et al. Geneious Basic: An integrated and extendable desktop software platform for the organization and analysis of sequence data. Bioinformatics. 2012;28: 1647–1649. doi:10.1093/bioinformatics/bts199

83. Poppe S, Dorsheimer L, Happel P, Stukenbrock EH. Rapidly Evolving Genes Are Key Players in Host Specialization and Virulence of the Fungal Wheat Pathogen Zymoseptoria tritici (Mycosphaerella graminicola). PLOS Pathog. 2015;11: 1–21.doi:10.1371/journal.ppat.1005055

84. Gibson DG, Young L, Chuang RY, Venter JC, Hutchison CA, Smith HO. Enzymatic assembly of DNA molecules up to several hundred kilobases. Nat Methods. 2009;6: 343–345. doi:10.1038/nmeth.1318

85. Krueger F, Andrews SR. Bismark: a flexible aligner and methylation caller for Bisulfite-Seq applications. Bioinformatics. 2011;27: 1571–1572. doi:10.1093/bioinformatics/btr167

86. Quinlan AR, Hall IM. BEDTools: a flexible suite of utilities for comparing genomic features. Bioinformatics. 2010;26: 841–842. doi.org/10.1093/bioinformatics/btq033

87. Southern EM. Detection of specific sequences among DNA fragments separated by gel electrophoresis. J Mol Biol. 1975;98: 503–517. doi:10.1016/S0022-2836(75)80083-0

88. Sambrock J, W. Russel D. Molecular Cloning: A Laboratory Manual(3rd edition). 2001

89. Marçais G, Kingsford C. A fast, lock-free approach for efficient parallel counting of occurrences of k-mers. Bioinformatics. 2011;27: 764–770. doi:10.1093/bioinformatics/btr011

90. R Core Team. R: A Language and Environment for Statistical Computing. R Found Stat Comput. 2019; Available: https://www.r-project.org

91. Lê S, Josse J, Husson F. FactoMineR: An R Package for Multivariate Analysis. J Stat Software. 2008, Vol 25, Issue 1; doi:10.18637/jss.v025.i01

92. Letunic I, Bork P. 20 years of the SMART protein domain annotation resource. Nucleic Acids Res. 2017;46: D493–D496. doi:10.1093/nar/gkx922

93. Finn RD, Attwood TK, Babbitt PC, Bateman A, Bork P, Bridge AJ, et al. InterPro in 2017-beyond protein family and domain annotations. Nucleic Acids Res. 2017;45: D190–D199. doi:10.1093/nar/gkw1107

