## Supplementary material for "Recent loss of the Dim2 DNA methyltransferase decreases mutation rate in repeats and changes evolutionary trajectory in a fungal pathogen": S1 Text

1) **Bisulfite data analysis pipeline**

**Quality control – Trimmomatic version 0.36 (1)**

*java -jar trimmomatic-0.36.jar PE -threads 8 -phred33 input_reads_R1.fastq.gz input_reads_R2.fastq.gz output_reads_paired.fastq output_reads_R1_unpaired.fastq output_reads_R2_paired.fastq output_reads_R2_unpaired.fastq HEADCROP:10 CROP:140 LEADING:3 TRAILING:3 SLIDINGWINDOW:4:15 MINLEN:50*

**Mapping and deduplication – Bismark version 0.20.0 (2)**

*perl bismark -n 1 path/to/genome -1 output_reads_paired_R1.fastq -2 output_reads_paired_R2.fastq*

*perl deduplicate_bismark --bam mapping_paired_bismark_bt2_pe.bam*

**Sorting and indexing – samtools version 1.5 (3)**

*samtools sort mapping_paired_bismark_bt2_pe.deduplicated.bam -o mapping_paired_bismark_bt2_pe.deduplicated_sorted.bam*

*samtools index mapping_paired_bismark_bt2_pe.deduplicated_sorted.bam*

**Extracting methylated sites – Bismark version 0.20.0 (2)**

*perl bismark_methylation_extractor -p --no_overlap --gzip --CX --bedgraph --cytosine_report --genome_folder path/to/genome -o output/directory mapping_paired_bismark_bt2_pe.deduplicated.bam*

**Separate CpG, CHG and CHH sites from methylation report generates with bismark_methylation_extractor**

zgrep "CHH" CX_report.txt.gz >> CHH_report.txt

zgrep "CHG" CX_report.txt.gz >> CHG_report.txt

zgrep -v "CCG" CX_report.txt.gz | grep "CG" >> CG_report.txt

**Filter methylated sites that are supported by at least 4 reads and more than 50 % of the mapped reads**

awk '{ if(($4 >= 1) && ($4 >= $5)) { print }}' CHG_report.txt > CHG_report_methylated.txt

awk '{ if($4 >= 4) { print }}' CHG_report_methylated.txt > CHG_report_methylated_4.txt

awk '{ if(($4 >= 1) && ($4 >= $5)) { print }}' CG_report.txt > CG_report_methylated.txt

awk '{ if($4 >= 4) { print }}' CG_report_methylated.txt > CG_report_methylated_4.txt

awk '{ if(($4 >= 1) && ($4 >= $5)) { print }}' CHH_report.txt > CHH_report_methylated.txt

awk '{ if($4 >= 4) { print }}' CHH_report_methylated.txt > CHH_report_methylated_4.txt

**Concatenate filtered methylation site files**

cat CHG_methylated_4.txt CG_methylated_4.txt CHH_methylated_4.txt > CX_methylated_4.txt

**Correlate methylated sites to genomic features – bedtools version 2.25.0 (4)**

bedtools intersect -wa -wb -a CX_methylated_4.bed -b gene/TE.bed > CX_intersect_gene/TE.txt

2) **Mapping of genome sequencing data and SNP analysis**

Paired-end reads of 250 bp were mapped directly to the genome of the reference isolate IPO323 (5) and Zt10(6). Processing of the reads was carried out using the below listed pipeline:

1) **Quality filtering using Trimmomatic version 0.38 (1)**

*java -jar /trimmomatic-0.38.jar PE R1.fastq R2.fastq R1_paired.fastq R1_unpaired.fastq R2_paired.fastq R2_unpaired.fastq ILLUMINACLIP:TruSeq3-PE.fa:2:30:10 LEADING:25 SLIDINGWINDOW:4:25 AVGQUAL:25 MINLEN:50*

2) **Mapping to reference genome using Bowtie 2 version 2.3.5 (7) and samtools version 1.7 (3)**

*bowtie2 --sensitive -x reference.fasta -1 R1_paired.fastq -2 R2_paired.fastq |*

*samtools view -q10 -bS |*

*samtools sort –o sorted.bam*

3) **Marking of duplicates, adding readgroup information and resorting using Picard version 2.18.20**

*java -jar picard.jar AddOrReplaceReadGroups INPUT= sorted.bam \*

*OUTPUT=sorted_RG.bam \*

*RGID=sample_ID RGLB=lib1 \*

*RGPL=ILLUMINA RGPU=unknown \*

*RGSM=sample_ID SORT_ORDER=coordinate*

*Java -jar picard.jar MarkDuplicates I= sorted_RG.bam \*

*O=sorted_RG_Dedup.bam M=sorted_RG_Dedup.txt*

4) **SNP calling using samtools (version 1.7) and bcftools (version 1.6)**

*samtools mpileup -E -C50 -Q20 -q20 -uf IPO323_reference R.bam | bcftools call --ploidy-file ploidy.txt -vc -O u -o raw.bcf*

5) **Filtering of SNPs using bcftools (version 1.6)**

*bcftools filter -o Q20DP8AF08.bcf -e'QUAL<20 | DP<8 | AF<0.8' raw.vcf*

6) **Pairwise comparison to progenitor strain to output SNPs unique to evolved strains using bcftools (version1.6)**

*bcftools isec -c all -p sample Q20DP8AF08.bcf.gz ref.fasta > private_to_progenitor.vcf*

1.	Bolger AM, Lohse M, Usadel B (2014) Trimmomatic: A flexible trimmer for Illumina sequence data. *Bioinformatics* 30(15):2114–2120.

2.	Krueger F, Andrews SR (2011) Bismark: a flexible aligner and methylation caller for Bisulfite-Seq applications. *Bioinformatics* 27(11):1571–1572.

3.	Li H (2011) A statistical framework for SNP calling, mutation discovery, association mapping and population genetical parameter estimation from sequencing data. *Bioinformatics* 27(21):2987–2993.

4.	Quinlan AR, Hall IM (2010) BEDTools: a flexible suite of utilities for comparing genomic features. *Bioinformatics* 26(6):841–842.

5.	Goodwin SB, et al. (2011) Finished Genome of the Fungal Wheat Pathogen Mycosphaerella graminicola Reveals Dispensome Structure, Chromosome Plasticity, and Stealth Pathogenesis. *PLoS Genet* 7(6):e1002070.

6.	Haueisen J, et al. (2019) Highly flexible infection programs in a specialized wheat pathogen. *Ecol Evol* 9(1):275–294.

7.	Langmead B, Salzberg SL (2012) Fast gapped-read alignment with Bowtie 2. *Nat Methods* 9(4):357–359.
